# MTFP1 preserves β-cell cristae structure and bioenergetics to ensure insulin release and glucose homeostasis

**DOI:** 10.64898/2026.01.12.697845

**Authors:** Sunehera Sarwat, Rosa Alén, Zhiyi Wu, Chuyue Zhang, Michael Paszek, MingMing Yang, Giada Ostinelli, Alina Mihalovits, Bell Baker, Timothy Wai, Guy A. Rutter, Tristan A. Rodriguez, Aida Martinez-Sanchez

## Abstract

Pancreatic β-cells are uniquely dependent on mitochondrial metabolism to couple glucose sensing to insulin secretion, a process impaired in diabetes.

Mitochondrial fission process 1 (MTFP1) is an inner mitochondrial membrane protein that plays pleiotropic, tissue-specific roles in mitochondrial function and dynamics. Our previous work has identified *Mtfp1* mRNA as a target for miR-125b, a microRNA that negatively regulates insulin secretion from β-cells. Nevertheless, the function of MTFP1 in these cells remained unexplored.

Here, we show that MTFP1 is essential for normal glucose-stimulated insulin secretion (GSIS) in mouse and human cell lines and islets, and that mice with β-cell-specific elimination of MTFP1 develop glucose intolerance. Whereas β-cell survival and mitochondrial content were unaffected, oxidative phosphorylation and ATP production were sharply lowered. These changes were accompanied by disruption of mitochondrial cristae structure and a reduced contact surface with the endoplasmic reticulum, providing a mechanistic basis for defective stimulus-secretion coupling. Conversely, MTFP1 overexpression in mouse and human islets sufficed to improve mitochondrial respiration and GSIS. Finally, MTFP1 downregulation blocked the positive effects of miR-125b elimination in GSIS and mitochondrial respiration, unveiling MTFP1 as a downstream effector of miR-125b.

Together, our findings identify MTFP1 as a critical regulator of β-cell mitochondrial architecture and function, necessary for efficient insulin secretion and glucose homeostasis, and a potential therapeutic target to enhance β-cell bioenergetic resilience in diabetes.

## Introduction

Mitochondria play a central role in the secretory function of pancreatic β-cells by coupling glucose metabolism to insulin secretion and by regulating stress responses and survival[1]. Once glucose enters the β-cell, it is rapidly metabolised via glycolysis and mitochondrial oxidative metabolism, leading to a sharp rise in the intracellular ATP/ADP ratio. This rise closes ATP-sensitive K+ (K_ATP_) channels, leading to plasma membrane depolarisation and subsequent Ca^2+^ influx into the cytosol, ultimately triggering insulin exocytosis[2].

Given the critical role of the organelle, mitochondrial dysfunction is increasingly recognised as a key driver of β-cell failure in both type 1 and type 2 diabetes (T1D; T2D). Islets from T2D donors exhibit impaired glucose stimulated insulin secretion (GSIS), along with marked defects in mitochondrial respiration[3] ultrastructure[4] and the expression of genes involved in mitochondrial function[5]. In T1D, mitochondrial abnormalities have also been linked to chronic β-cell inflammation and increased susceptibility to immune attack[6]. More recently, altered mitochondrial activity has been implicated in the reduced functionality of induced pluripotent stem cell–derived islets compared to primary human islets, thereby limiting their ability to restore glucose control after transplantation and their therapeutic application[7, 8]. Furthermore, commonly used anti-hyperglycemic drugs such as metformin and G-protein coupled receptors (GPCRs) agonists, including those for glucagon-like peptide-1 (GLP-1) may exert their effects on β-cells in part by modulating mitochondrial dynamics and function [9, 10]. Nevertheless, the cellular and molecular mechanisms that maintain mitochondrial fitness in healthy and diseased β-cells remain poorly understood, limiting the development of effective mitochondria-targeted therapies for diabetes.

Mitochondrial fusion and fission are highly dynamic, tightly balanced processes essential for mitochondrial quality control and adequate function[11]. The roles of several regulatory proteins of the fusion and fission machinery have been characterised in β-cells, including the fusion proteins, GTPases MFN1/2 (mitofusins 1 and -2) and OPA1 (optic atrophy protein) and the fission proteins FIS1 (transmembrane fission protein 1), MFF (mitochondrial fission factor) and the GTPase dynamin-related protein 1 (DRP1)[1]. It is, however, debated whether mitochondrial fragmentation promotes apoptosis or correlates with hyperglycaemia or diabetes[11]. For example, the elimination of *Mfn1/2* results in mitochondrial fragmentation and impairs mitochondrial and secretory function[12], whilst deletion of *Drp1* results in enlarged mitochondria and impaired insulin secretion[13]. Furthermore, increased mitochondrial fragmentation has been observed in murine models of diabetes[14] and mitochondrial structural defects correlate with mitochondrial dysfunction in human T2D donors[4]. It is, however, unclear whether these changes contribute to the development of the disease and/or are secondary to defective mitochondrial dynamics caused by glucolipotoxicity[15].

Mitochondrial fission process 1 (MTFP1), also known as mitochondrial protein 18KDa (MTP18), encoded by the *MTFP1* gene, is a protein localised to the mitochondrial inner membrane (IMM) and initially described to promote mitochondrial fission[16, 17]. Later studies unveiled a more complex role for MTFP1 in the regulation of mitochondrial and cellular function, which differs markedly between cell types/tissues and between *in vitro* or *in vivo* settings. For example, MTFP1 promotes mitochondrial fragmentation and increases survival and growth of several cancer cell lines[18–20], whilst it mediates apoptosis in rat proximal tubular cells[21]. Strikingly, cardiomyocyte-specific deletion of *Mtfp1* in mice resulted in lethal adult-onset dilated cardiomyopathy due to defective IMM proton leak and increased sensitivity to mitochondrial permeability transition pore (mPTP) opening in the absence of apparent changes in mitochondrial fragmentation[22]. Remarkably, the same researchers observed that hepatocyte-specific *Mtfp1* ablation protected mice against high-fat diet-induced steatosis[23]. Hepatocytes from these mice showed increased oxidative phosphorylation (OXPHOS) capacity and mitochondrial respiration, as well as lowered mPTP opening and subsequent protection against apoptotic liver damage, also in the absence of changes in mitochondrial morphology[23]. Recently, Tabara et al.[24] uncovered a novel mechanism underlying the impact of MTFP1 in mitochondrial morphology, suggesting that MTFP1 functions as a negative regulator of IMM fusion rather than promoting fission, at least *in vitro*. Accordingly, in U2OS cells, MTFP1 prevents the fusion of damaged mitochondria, which are then degraded via autophagosomes to maintain adequate mtDNA content [24]. Together, these observations underscore the highly context-dependent functions of MTFP1, highlighting the need to examine its role in a cell-specific manner. Despite this emerging complexity, virtually nothing is known about the role of MTFP1 in β-cells nor how its expression is regulated.

Using an unbiased high-throughput approach, our lab previously identified *MTFP1* as a direct gene target of microRNA (miRNA) miR-125b in pancreatic β-cells. MiR-125b is a negative regulator of β-cell function, whose expression increases in response to hyperglycaemia[25, 26]. MiR-125b overexpression *in vitro* and *in vivo* impaired insulin secretion [25] whereas elimination of miR-125b in human pancreatic EndoCβ-H1 cells (EndoCβ-H1-MIR125B-KO) resulted in enhanced glucose-stimulated insulin secretion and shortened mitochondria [25]. Whilst the expression of genes with well-characterised roles in β-cell mitochondrial fission and fusion remained unchanged, EndoCβH1-MIR125B2-KO cells contained higher levels of *MTFP1* mRNA [25]. These findings suggested that miR-125b may regulate β-cell mitochondrial behaviour and secretory capacity at least in part through MTFP1, and led us to hypothesise that, in these cells, MTFP1 promotes secretory function by facilitating mitochondrial fission.

In the present study, we demonstrate that lowering MTFP1 expression in human β-cell lines and primary islets impairs glucose-stimulated insulin secretion (GSIS) and that, conversely, MTFP1 overexpression increases insulin secretion. These effects occur alongside changes in mitochondrial elongation and respiration, but in the absence of detectable changes in mtDNA content. Furthermore, we show that *MTFP1* is a major contributor to the positive effects of miR-125b elimination in insulin secretion from EndoCβH1 cells. Thus, knockdown of *MTFP1* in EndoCβH1-MIR125B2-KO cells prevents the improvement of GSIS caused by miR-125b deletion, as well as the miR-125b-mediated effects in mitochondrial shape and respiration.

To assess the role of β-cell MTFP1 *in* vivo, we generated β-cell specific knockout mice (MTFP1-βKO). Consistent with our findings *in* vitro, MTFP1-βKO mice developed glucose intolerance due to defective secretory function, in the absence of changes in β-cell survival and proliferation. Mechanistically, studies using isolated islets from these mice demonstrated that MTFP1 elimination strongly impaired ATP and intracellular Ca^2+^ rises that promote insulin secretion in response to exposure to high glucose concentrations. These effects reflected the strong reduction in OXPHOS and mitochondrial membrane potential observed in these islets. Surprisingly, MTFP1 elimination only had a modest effect on mitochondrial elongation in MTFP1-βKO or human islets, contrary to what we observed in β-cell lines, whereas mitochondrial DNA content remained unchanged in all the models. Importantly, electron micrographs of these islets unveiled defective cristae ultrastructure and a marked reduction in the contact surface with the endoplasmic reticulum (ER), consistent with metabolic uncoupling in the absence of detectable changes in OXPHOS gene expression.

Together, our findings identify MTFP1 as an essential regulator of β-cell mitochondrial architecture and bioenergetic function, required for efficient insulin secretion and glucose homeostasis.

## Results

### MTFP1 is essential for adequate glucose stimulated insulin secretion in EndoC-βH3 cells and human islets

To determine whether MTFP1 plays a role in GSIS in human β-cells we used CRISPR/CAS9 to knockout *MTFP1* in the human β-cell line EndoC-βH3[27]. Using two different pairs of gRNAs targeting exons 1-2 of *MTFP1*, we obtained two populations of EndoC-βH3 cells (EndoCβH3-MTFP1-KO1 and EndoCβH3-MTFP1-KO2) with > 40% reduction in *MTFP1* mRNA content (Figure 1A), despite not being able to detect the effect at the protein level, as MTFP1 remained under the detection level by Western blot in these cells.

**Figure 1.**
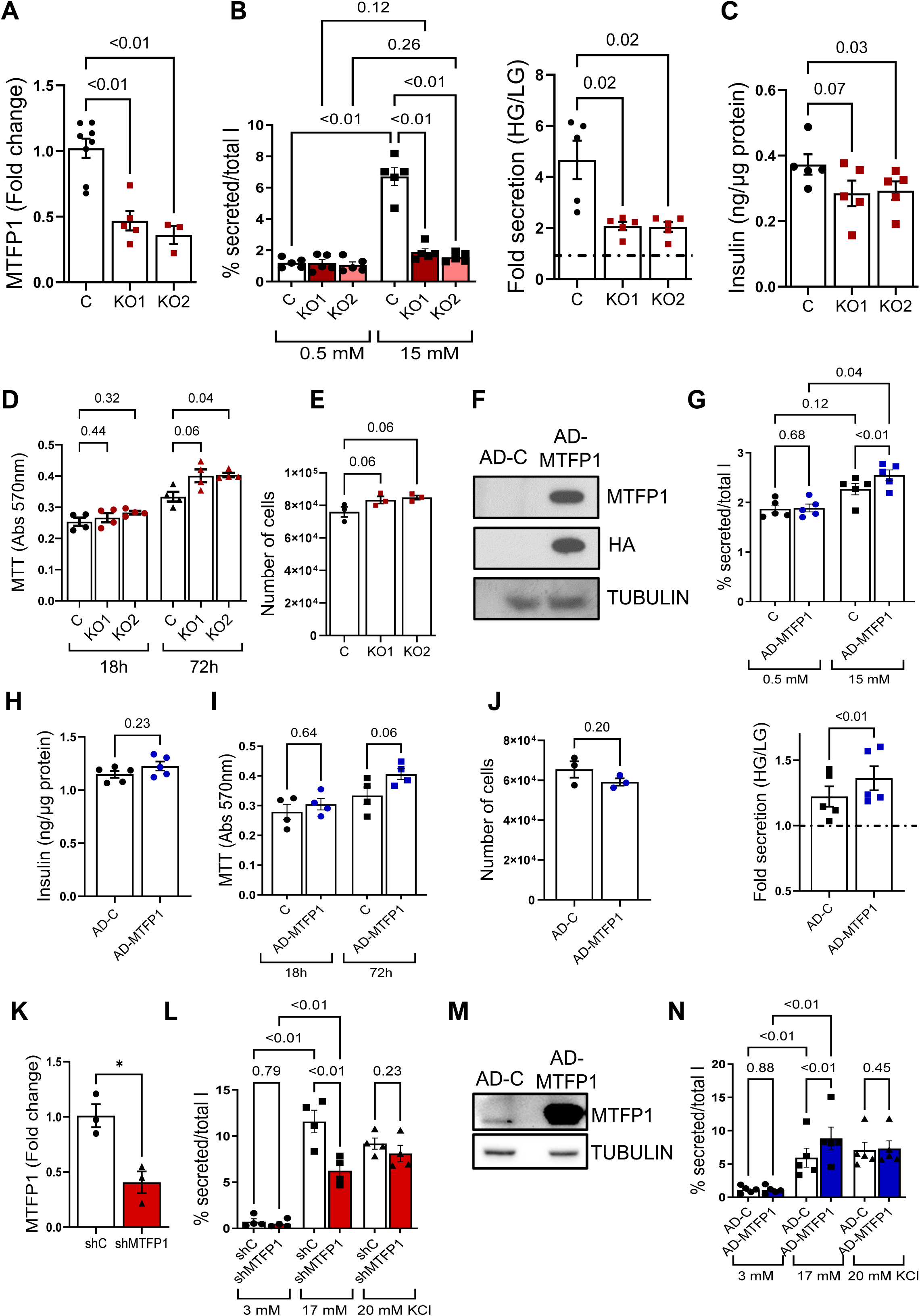
Loss- and gain-of MTFP1 function results in reduced and increased GSIS, respectively, in human β-cells. **A-E)** MTFP1-KO1 (KO1) and MTFP1-KO2 (KO2) EndoC-βH3 cells were generated with CRISPR/Cas9 targeting MTFP1. CRISPR/Cas9 EndoCβ-H3 populations were obtained via three independent lentiviral infections. **F-J)** EndoCβ-H3 cells were infected with adenovirus expressing HA-tagged MTFP1 and GFP (AD-MTFP1) or GFP only (AD-C, controls) at 2 MOI for 48h preceding the experiments. Cells were treated with tamoxifen for 21-42 days to enhance secretory function. **K-N)** Human islets were infected with lentivirus carrying an shRNA targeting MTFP1 (sh-MTFP1, KD) or non-targeting (sh-C, C) at 5 MOI **(K, L)** or with adenovirus expressing HA-tagged MTFP1 and GFP (AD-MTFP1) or GFP only (AD-C, controls) at 10 MOI **(M,N)** for 72h. **A)** RT-qPCR data showing expression of MTFP1. Data is normalized to RPLP0 and presented as fold-change from control (C). **B,G)** Glucose stimulated insulin secretion presented as % of total insulin or fold change (right-hand side panel). HG=15mM, LG=0.5mM glucose. **C,H)** Cellular insulin content as quantified with an HTRF and normalized to total protein content quantified with BCA. **D,I)** Cellular proliferation measured by MTT ((3-(4,5-dimethylthiazol-2-yl)-2,5-diphenyltetrazolium bromide) assay following 18h and 72h of equal cell numbers plating. **E,J)** Cell number quantified by counting of DAPI-negative cells 72h after plating of equal number of cells (6x10^4^). **F)** Western (immuno-) blot with antibodies against MTFP1 (MTFP1), the HA-tag (HA) or against tubulin (Tubulin), as housekeeping loading control. **K)** RT-qPCR data showing expression of MTFP1. Data is normalized to RPLP0 and presented as fold-change from control (C). **L,N)** Glucose stimulated insulin secretion presented as % of total insulin in response to high glucose concentration or KCl. HG=15mM, LG=0.5mM glucose. **M)** Western (immuno-) blot with antibodies against MTFP1 (MTFP1) and Tubulin, as housekeeping loading control. Each dot represents an independent experiment and, for human islets, separate donor (n=3-5). Error bars represent SEM, one-way ANOVA (A), one-way ANOVA with repeated measures (B-Fold secretion, C, E) and Dunnet multiple comparisons test, two-way ANOVA (repeated measures) and Tukey’s multiple comparisons test (B, D,G,I, L, N), and paired Student’s t test (H, G -Fold secretion, J, K).

Both EndoCβH3-MTFP1-KO1 and EndoCβH3-MTFP1-KO2 cells displayed a sharp lowering in insulin secretion in response to high glucose in comparison with control cells expressing non-targeting gRNAs (Figure 1B). These cells also showed a decrease in total insulin content (Figure 1C), whilst the number of metabolically active cells was marginally increased, as measured by MTT assays (Figure 1D) and total cell count (Figure 1E).

To determine whether MTFP1 overexpression suffices to improve GSIS, we infected EndoC-βH3 cells with an adenovirus expressing HA-tagged MTFP1 (Figure 1F). GSIS was increased in MTFP1-overexpressing EndoC-βH3 (EndoCβH3-MTFP1-OE) cells in comparison to cells infected with control adenovirus(Figure 1G), in the absence of significant differences in insulin content (Figure 1H) or growth (Figure 1I,J).

Given that EndoC-βH3 cells exhibit features of β-cell dedifferentiation and impaired secretory function compared with primary human islets, we next examined whether MTFP1 plays a similar role in primary human islets, where mature β-cell architecture and stimulus–secretion coupling are preserved. Accordingly, we infected human primary islets from non-diabetic cadaveric donors with MTFP1 shRNA-expressing lentivirus, resulting in a ∼60% reduction in *MTFP1* mRNA and a ∼50% reduction in insulin secretion in response to high glucose levels (Figure 1K, L) in comparison with islets transduced with non-targeting shRNAs. Conversely, human islets infected with virus overexpressing MTFP1 (AD-MTFP1) showed increased GSIS relative to GFP-only expressing controls (Figure 1M, N).

Overall, these data suggest that MTFP1 is essential and sufficient to stimulate insulin secretion in response to glucose in a human β-cells.

### MTFP1 regulates mitochondria respiration in EndoCβH3 cells

Given the previously proposed, cell type-dependent role of MTFP1 in mitochondrial dynamics, we next sought to investigate whether MTFP1 modulation affected mitochondrial morphology and mitochondrial respiration in EndoC-βH3 cells after MTFP1 gain or loss-of-function.

To this end, we imaged EndoCβH3-MTFP1-KO1/2 and EndoCβH3-MTFP1-OE cells, together with their corresponding controls, following incubation with the fluorescent dye Mitotracker. Contrary to our expectations, mitochondria elongation, circularity and perimeter remained comparable in cells with reduced MTFP1 expression and in control cells (Supplemental Figure 1). Likewise, neither the total number of mitochondria per cell nor the total mitochondrial area was altered (Supplemental Figure 1). Mitochondria morphology and area also remained unchanged in cells overexpressing MTFP1 compared with GFP-only controls (Supplemental Figure 1B). Consistent with these findings, mitotracker staining of human islets revealed a similar lack of effect of MTFP1 loss- or gain-of-function on mitochondrial morphology, with MTFP1 overexpression resulting only in a modest, non-significant trend towards shorter mitochondria (Supplemental Figure C1). Together, these data indicate that, in human β-cells, robust modulation of secretory function by MTFP1 occurs in the absence of detectable changes in mitochondrial length.

To examine the respiratory capacity of the EndoC-βH3 mutant cells, we subjected them to a mitochondrial stress test using a Seahorse^TM^ analyser. We sequentially exposed the cells to oligomycin -to inhibit ATP synthase and assess ATP-linked respiration, Carbonyl cyanide-*p*-trifluoromethoxyphenylhydrazone (FCCP) - to uncouple the mitochondrial membrane and determine maximal respiratory capacity, and rotenone/antimycin A - to block complexes I and III and measure non-mitochondrial respiration. Notably, acute high-glucose stimulation did not elicit a measurable increase in OCR under our assay conditions, regardless of genotype, consistent with the known metabolic limitations of EndoC β-cell lines, which do not fully recapitulate the oxidative metabolic coupling of primary human β-cells. In line with a greater reliance on non-mitochondrial ATP-generating pathways, we observed an increase in extracellular acidification rate (ECAR) upon high-glucose addition in these cells (data not shown).

Interestingly, EndoCβH3-MTFP1-KO1 cells showed significantly lowered basal respiration, ATP-linked respiration, maximal respiration and, unexpectedly, proton leak, whereas spare respiratory capacity remained largely unchanged (Figure 2A). Non-mitochondrial respiration was slightly, albeit significantly, reduced as well (Figure 2A). These data indicate that loss of MTFP1 compromises overall mitochondrial oxidative phosphorylation efficiency and lowers ATP production capacity, reflecting impaired mitochondrial function in EndoCβH3 cells. A similar analysis in EndoCβH3-MTFP1-OE cells revealed the opposite effect, with MTFP1 overexpression resulting in increased basal, ATP-linked maximal respiration and proton leak, without significant changes in spare respiratory capacity or non-mitochondrial respiration (Figure 2B).

**Figure 2.**
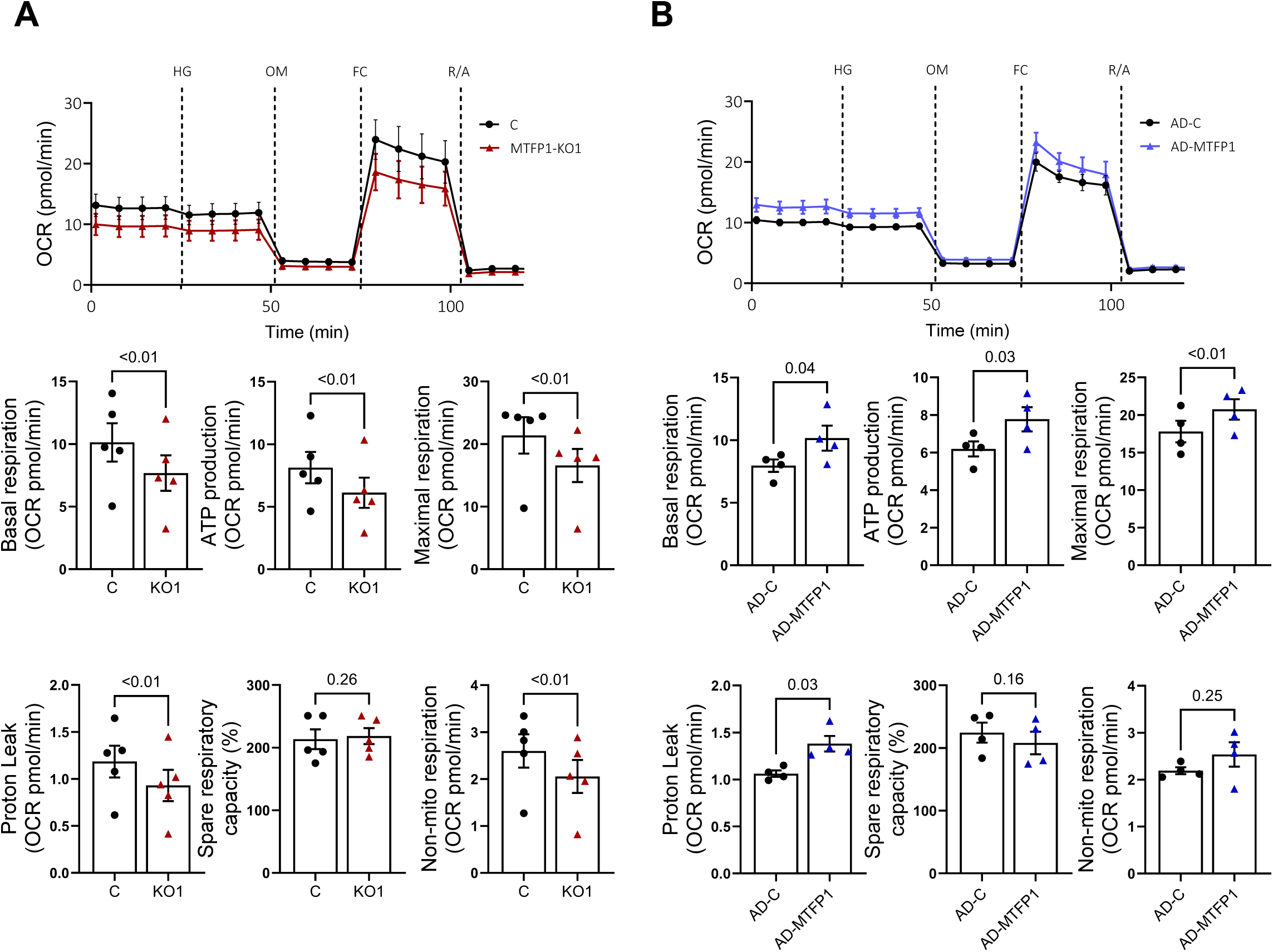
Loss- and gain-of MTFP1 function results in reduced and increased mitochondrial respiration in EndoCβ-H3 cells, respectively without affecting mtDNA content. **A)** MTFP1-KO1 (KO1) cells were generated with CRISPR/Cas9 targeting MTFP1. CRISPR/Cas9 EndoCβ-H3 populations were obtained via three independent lentiviral infections. **B)** EndoCβ-H3 cells were infected with adenovirus expressing HA-tagged MTFP1 and GFP (AD-MTFP1) or GFP only (AD-C, controls). EndoCβ-H3 were treated with tamoxifen for 21-42 days. Oxygen consumption rate (OCR) was quantified in a Seahorse analyzer upon addition of glucose (15 mM), oligomycin (2µM), FCCP (2µM) and rotenone and antimycin RotA/AA (2.5µM each). Parameters calculated are Basal respiration (OCR at 0.5 mM glucose), ATP production (Last rate before oligomycin injection – minimum rate after oligomycin injection), maximal respiration (Maximum FCCP rate – Non-Mitochondrial OCR), proton leak (Minimum rate after Oligomycin injection-Non-Mitochondrial OCR), spare respiratory capacity (Maximal over Basal Respiration, in %) and non-mitochondrial respiration (OCR following RotA/AA treatment). Each dot represents an independent experiment (n=5). Error bars represent SEM, two-way ANOVA (repeated measures) and Sidak’s multiple comparisons test (top graphs), and paired Student’s t test.

In osteosarcoma-derived U2OS cells, MTFP1 overexpression and deletion have been shown to reduce and increase, respectively, mitochondrial DNA (mtDNA) content[24]. As altered mtDNA content in β-cells has been extensively shown to trigger mitochondrial and cellular dysfunction even in the face of maintained mitochondrial area[28, 29], we investigated whether MTFP1 modulation resulted in changes in mtDNA. However, consistent with the absence of significant changes in mitochondrial area per cell, mtDNA content remained unchanged in MTFP1-overexpressing cells and was slightly reduced, rather than increased, in MTFP1-deficient cells compared with controls. These findings indicate that the bioenergetic effects of MTFP1 modulation occur largely independently of mtDNA copy number (Supplemental Figure 2).

### MTFP1 is a key mediator of the improved oxidative and secretory capacity observed in miR-125b–deficient EndoC-βH1 cells

We have previously shown that miR-125b elimination from human EndoC-βH1 cells (EndoCβ-H1-MIR125B2-KO) results in mitochondrial shortening and improved GSIS, as well as increased expression of its target *MTFP1*[25]. To investigate whether these effects may be mediated by MTFP1, we silenced MTFP1 using shRNA-expressing lentivirus in EndoCβH1-MIR125B2-KO cells and assessed mitochondrial elongation and insulin secretion. As previously observed, MTFP1 expression increased following miR-125b elimination; however, this effect was prevented by the presence of MTFP1 shRNAs, which effectively reduced MTFP1 mRNA levels in EndoCβ-H1-MIR125B2-KO cells and, to a lesser extent, in control cells (Figure 3A). MTFP1 downregulation completely abolished the improvement in GSIS mediated by miR-125b elimination (Figure 3B). Moreover, MTFP1 knockdown fully reversed the mitochondrial shortening observed in EndoCβH1-MIR125B2-KO cells (Figure 3C), despite having no detectable effect in EndoCβH1-Control cells or following its deletion in EndoCβH3 cells (EndoCβH3-MTFP1-KO; Supplemental Figure 1).

**Figure 3.**
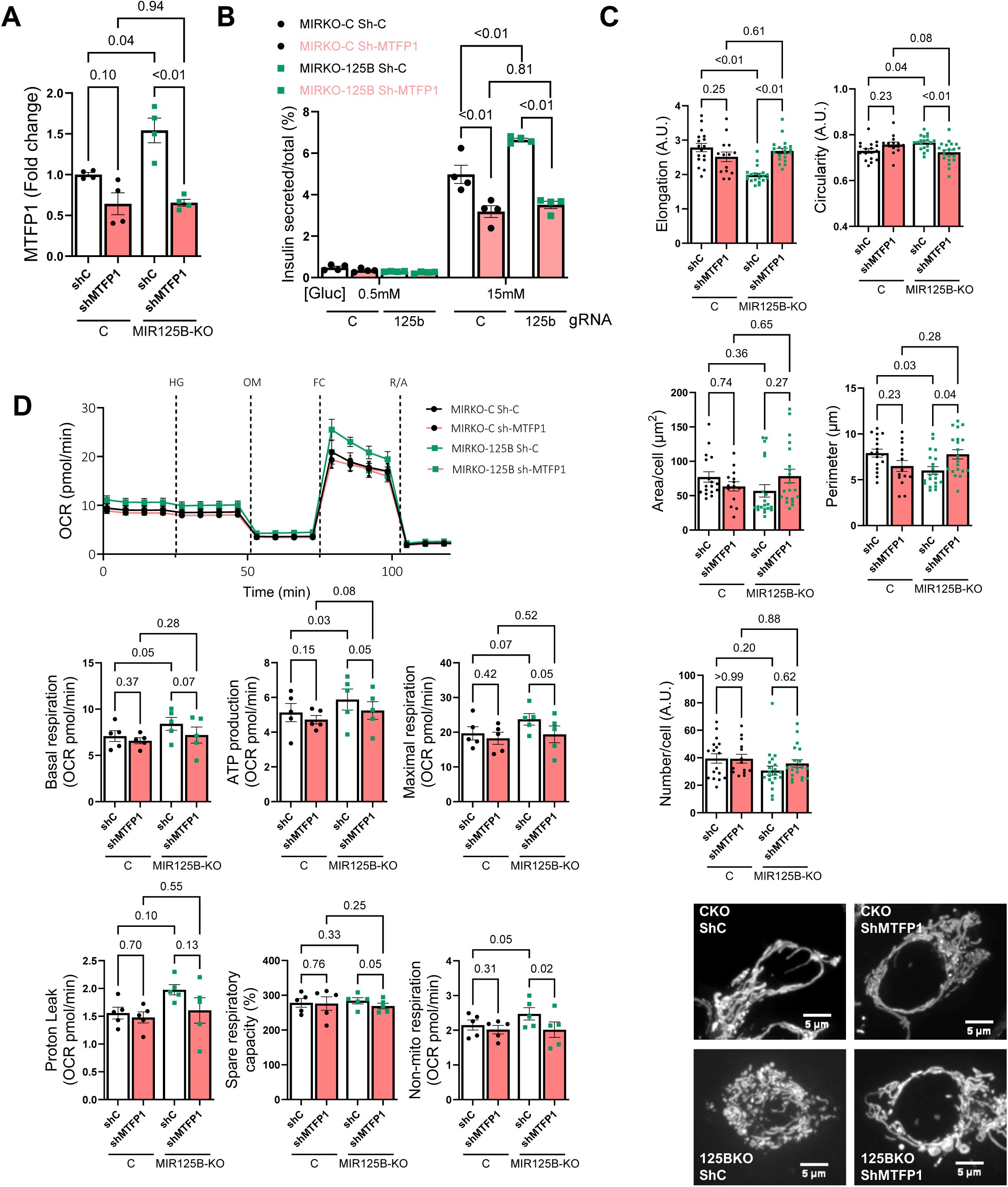
MTFP1 mediates some of the positive effects of miR-125b elimination in insulin secretion and respiration. Cells containing CRISPR-Cas9 alone (Control, C, MIRKO-C) or with gRNAs targeting pre-miR-125b-2 (EndoCβ-H1-MIR125B-KO, KO, MIRKO-125B) were infected with lentivirus expressing shRNAs targeting *MTFP1* (shMTFP1) or non-targeting (shC) for 48h. **A)** RT-qPCR data showing expression of MTFP1. Data is normalized to RPLP0 and presented as fold-change from control. **B)** Glucose stimulated insulin secretion presented as % of total insulin in response to high glucose concentration. **C)** Mitochondrial network was stained with mitotracker-green in live cells. ImageJ was used to quantify individual mitochondrial length, circularity, number of mitochondrial per islet area imaged, and total mitochondria per islet area imaged. Representative images are shown on the bottom. **D)** Oxygen consumption rate (OCR) was quantified in a Seahorse analyzer upon addition of glucose (15 mM), oligomycin (2µM), FCCP (2µM) and rotenone and antimycin RotA/AA (2.5µM each). Parameters calculated are Basal respiration (OCR at 0.5 mM glucose), ATP production (Last rate before oligomycin injection – minimum rate after oligomycin injection), maximal respiration (Maximum FCCP rate – Non-Mitochondrial OCR), proton leak (Minimum rate after Oligomycin injection-Non-Mitochondrial OCR), spare respiratory capacity (Maximal over Basal Respiration, in %) and non-mitochondrial respiration (OCR following RotA/AA treatment). Each dot represents an independent experiment (A-B,D, n=3-5) or one image acquisition ((C), n= 4-10 images/experiment, n=3 experiments). Error bars represent SEM, two-way ANOVA (A-D; repeated measures (A, D)) with Fisher’s LSD (A, D), with Tukey’s multiple comparisons test (C), and three-way ANOVA (repeated measures) with Tukey’s multiple comparisons test (B).

Although our previous work did not investigate the effect of miR-125b elimination on EndoCβ-H1 cells’ respiration, Seahorse experiments demonstrated that EndoCβ-H1-MIR125B2-KO cells displayed enhanced mitochondrial respiration, which was lost following MTFP1 knockdown, with basal, ATP-linked, and maximal respiration returning to control levels (Figure 3D).

Together, these results demonstrate that MTFP1 upregulation is required for the beneficial effects of miR-125b deletion on β-cell mitochondrial and secretory function.

### **β**-cell MTFP1 is essential for adequate insulin secretion and glucose homeostasis *in vivo*

To investigate the role of β-cell MTFP1 *in vivo* and determine whether its capacity to control insulin secretion in cell lines results in physiologically relevant effects on whole body glucose control, we generated mice with β-cell specific deletion of MTFP1. To do so, we bred *MTFP1-*floxed mice[22] with the highly selective Ins1-Cre line[30] generating MTFP1-βKO mice (*Mtfp1*^lox/lox^, Ins-Cre^+^) and littermate controls (*Mtfp1*^lox/lox^). Notably, a single Ins1-Cre allele has been extensively demonstrated by us and others not to affect glucose tolerance, insulin content, or β-cell function[30–32]. MTFP1 elimination was confirmed in isolated islets from MTFP1-βKO mice, where the near-complete loss of the protein indicates that the bulk of MTFP1 (protein) expression in mouse islets derives from β-cells (Figure 4A).

**Figure 4.**
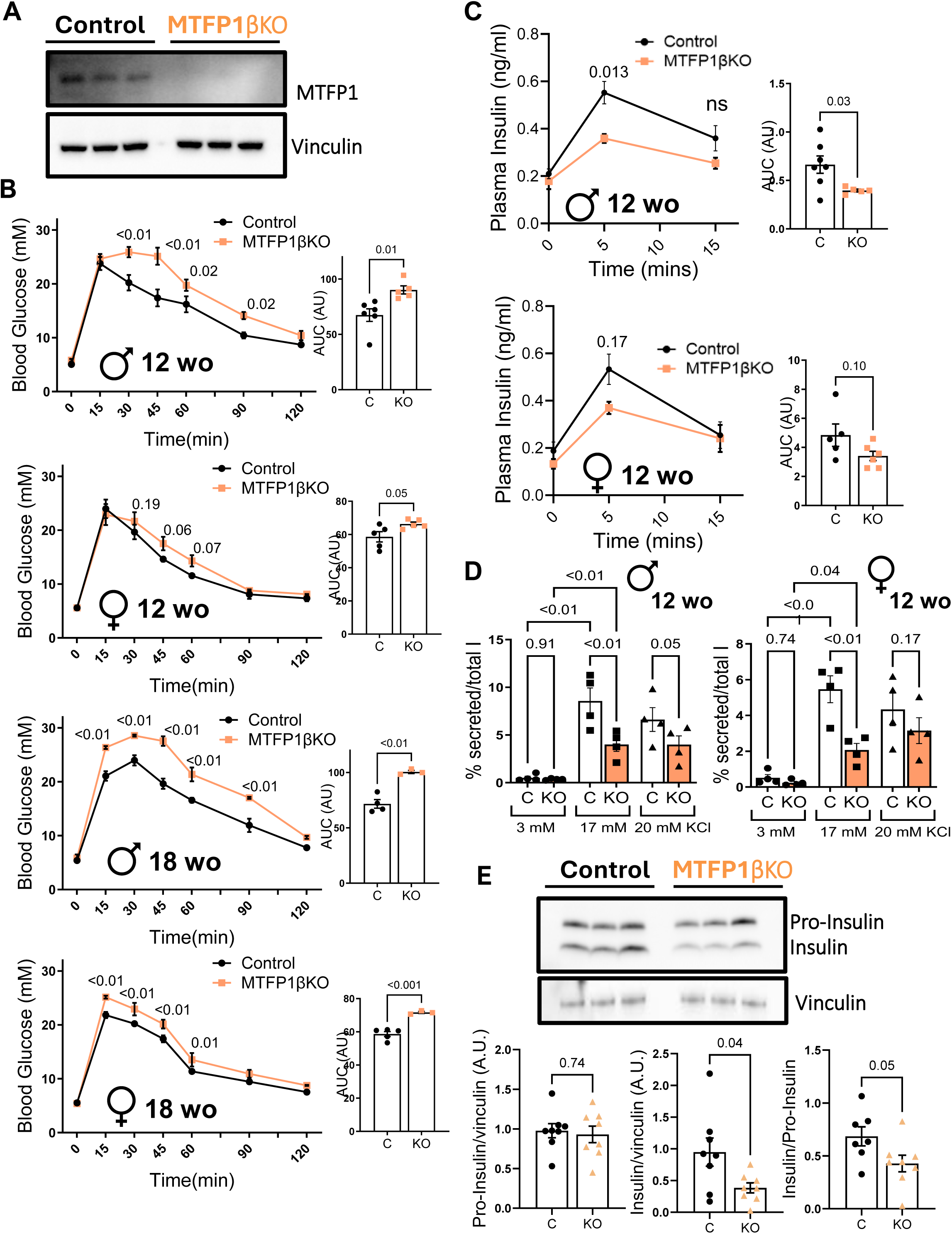
β-cell-selective elimination of (MTFP1βKO)results in glucose intolerance and impaired glucose stimulated insulin secretion. **A)** Representative Western blot (WB) showing MTFP1 expression in isolated islets from control (MTFP1^f/f^) and MTFP1βKO mice (MTFP1^f/f^, InsCre^+/−^). **B)** Glucose tolerance test in 12- and 18-week-old male (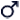) and female (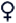) MTFP1βKO and littermate control mice. **C)** Insulin secretion, induced by 3 g/kg glucose, was assessed in 12-week-old male (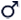) and female (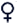) MTFP1βKO and littermate control mice. Area under the curves (AUC) are presented on the right-hand side of each graph. **D)** Insulin secretion of isolated islets in response to glucose and KCl. **E)** Representative Western blot (WB) showing Insulin and Pro-Insulin expression in isolated islets. Bar graphs show densitometry quantification normalized to vinculin. Each dot represents a different mouse, n=3-8 mice/genotype. AU = arbitrary units. Error bars represent SEM, two-way ANOVA (repeated measures) and Fisher’s LSD (B, C), Student’s t test (AUC B,C) and Welch’s t test (E).

Both male and female MTFP1-βKO mice had comparable weight and fed glycaemia measured up to 18 weeks of age (Supplemental Figure 3A, B). Nevertheless, knockout mice displayed strong glucose intolerance as measured during intraperitoneal glucose tolerance tests (IPGTT) from 12 and 18 weeks of age in males and females, respectively (Figure 4B), whereas insulin tolerance remained largely unaffected (Supplemental Figure 3C). Consistent with underlying β-cell defects, insulin secretion in response to a bolus of glucose administered intraperitoneally was also reduced in 12-week-old male and, to a lesser extent, female mice (Figure 4C). These defects occurred in the absence of detectable changes in β-cell mass, growth or apoptosis (Supplemental Figure 3D), pointing to a metabolic regulation of β-cell function by MTFP1.

Confirming an islet autonomous effect, GSIS was strongly reduced in both male and female MTFP1-βKO isolated islets in comparison to controls (Figure 4D), whereas secretion following KCl-mediated depolarisation of the plasma membrane was not significantly reduced. Western blot of the isolated islets revealed a reduction in their mature insulin content, though pro-insulin remained unchanged (Figure 4E), indicating impaired proinsulin processing.

These findings overall demonstrate that MTFP1 is essential to maintain insulin production and secretion in β-cells and, consequently, glucose homeostasis in both male and female mice *in vivo*.

### β-cell MTFP1 is essential to maintain mitochondrial membrane potential and glucose stimulated ATP generation

In β-cells, glucose uptake results in enhanced glycolytic pyruvate production. Pyruvate then enters the mitochondria where it serves as a substrate for the tricarboxylic acid (TCA) cycle and drives oxidative phosphorylation, leading to increased ATP generation. The resulting rise in the cytosolic ATP/ADP ratio closes ATP-sensitive K⁺ (K_ATP_) channels, causing membrane depolarisation, Ca²⁺ influx, and ultimately insulin secretion[2]. When quantified in isolated islets transduced with adenovirus encoding the ATP sensor Perceval[33] we observed a sharp reduction in the initial glucose-induced increases in cytosolic ATP in MTFP1-βKO islets (Figure 5A).

**Figure 5.**
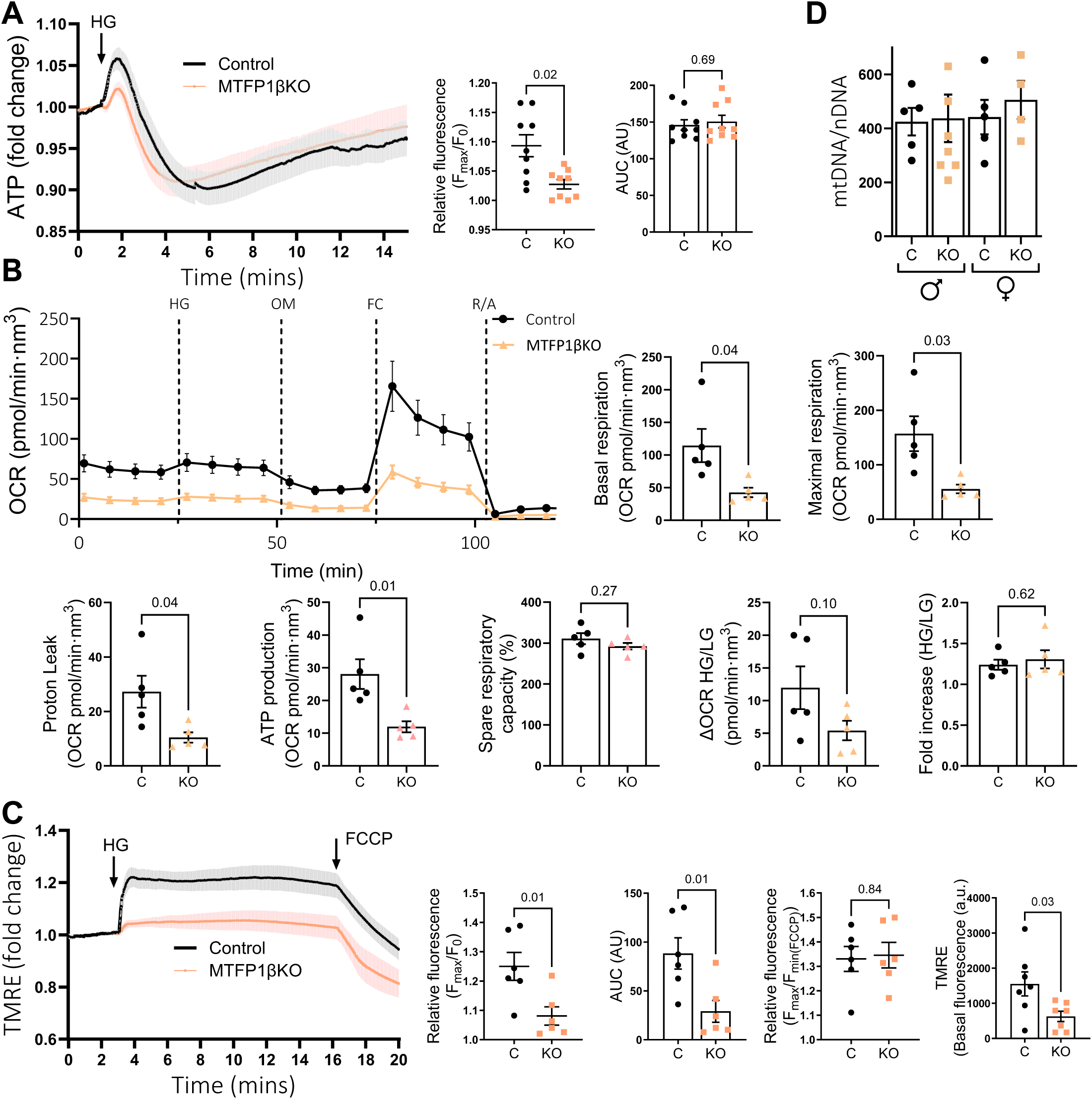
MTFP1-βKO islets show defects in mitochondrial respiration and their capacity to produce ATP. Islets were isolated from 12-week old male control and MTFP1βKO mice. **A)** ATP production in response to high glucose (HG=17 mM) measured upon 48h incubation with fluorescence Perceval probe and presented as fold change from basal. **B)** Oxygen consumption rate (OCR) measured upon addition of high glucose (17 mM), oligomycin (2µM), FCCP (2µM) and rotenone and antimycin RotA/AA (2.5µM each). Parameters calculated are Basal respiration (OCR at 0.5 mM glucose), ATP production (Last rate before oligomycin injection – minimum rate after oligomycin injection), maximal respiration (Maximum FCCP rate – Non-Mitochondrial OCR), proton leak (Minimum rate after Oligomycin injection-Non-Mitochondrial OCR) and spare respiratory capacity (Maximal over Basal Respiration, in %). **C)** Mitochondrial membrane potential measured upon incubation with TMRE in response to high glucose (HG=17 mM) and FCCP (1 μm) presented as fold change from basal or raw signal (Basal fluorescence, right-hand-side panel, AU= arbitrary units). **D)** MtDNA copy number was calculated as the ratio of the mitochondrial encoded gene *mt-ND1* to the nuclear *HK1*. For A-C, all islets were initially incubated at 3mM glucose. Each dot represents a single image acquisition (each including 3-8 islets) from 3-4 different mice/genotype (A,B,C) or islets from a single mouse (n=5/genotype), averaged from technical replicates (B,D). Error bars represent SEM, Student’s t test (A-D) and Welch’s t test (B).

Similar to our observations in EndoCβH3-MTFP1-KO cells, oxygen consumption rate (OCR) profiles in isolated MTFP1-βKO and control islets revealed a strong reduction in basal and maximal respiration, whereas their spare respiratory capacity remained unchanged (Figure 5B).

Consistent with our previous measurements using Perceval, MTFP1-βKO islets also displayed a marked reduction in ATP-linked respiration, as well as in the increase in OCR observed in response to high-glucose exposure, albeit this did not reach statistical significance (Figure 5B).

Interestingly, analysis of mitochondrial membrane potential using tetramethyl rhodamine ethyl ester (TMRE) showed a strong lowering of both basal and high glucose-induced membrane potential in MTFP1-βKO islets, whilst FCCP treatment effectively collapsed the membrane potential in both control and MTFP1-βKO islets (Figure 5C).

Importantly, and consistent with our findings in EndoCβH3 with MTFP1-gain/loss-of-function, mtDNA content was unchanged between control and MTFP1-βKO islets, indicating that the bioenergetic defects arise independently of mtDNA copy number (Figure 5D).

Overall, a concurrent reduction in mitochondria membrane potential and oxygen consumption in MTFP1-deficient β-cells suggests a defect in proton motive generation rather than altered coupling efficiency.

### **β**-cell MTFP1 is essential to maintain mitochondrial cristae structure

Reduced expression of OXPHOS components could be a plausible explanation for the defects observed in OCR and ATP production[34]. Accordingly, we performed Western immunoblotting of MTFP1-βKO and control islets using a cocktail of antibodies against OXPHOS complexes I to V. As shown in Supplemental Figure 4A, no significant changes were observed in the levels of NDUFB8 (CI), SDHB (CII), UQCRC2 (CIII), MTCO1 (CIV) and ATP5A (CV) in MTFP1-βKO islets. The expression of these proteins also remained unchanged in EndoCβH3-MTFP1-KO and EndoCβH3-MTFP1-OE cells (Supplemental Figure 4B).

Another plausible explanation for the bioenergetic defects observed upon MTFP1 deletion could be that loss of the protein alters mitochondrial ultrastructure, compromising cristae organization and efficiency of oxidative phosphorylation without affecting overall ETC protein abundance. In support of this possibility, transmission electron microscopy (TEM) revealed that MTFP1-βKO mitochondria in MTFP1-βKO β-cells have a lower proportion of normal cristae, blindly assigned using a cristae score (Figure 6A, B). Furthermore, quantitative morphometric analysis of MTFP1-βKO β-cells mitochondria unveiled a significant decrease in cristae volume per mitochondria (Figure 6C), defined as the percentage of cristae area occupying the mitochondrial area. At the whole-organelle level, the total number of mitochondria per β-cell area imaged remained comparable between genotypes (Figure 6D), although the total mitochondrial area fraction was increased (Figure 6E). This was a result from MTFP1-βKO mitochondria being more elongated and branched compared to controls, as quantitatively reflected by a significant increase in form factor and lower circularity (Figure 6F, G).

**Figure 6.**
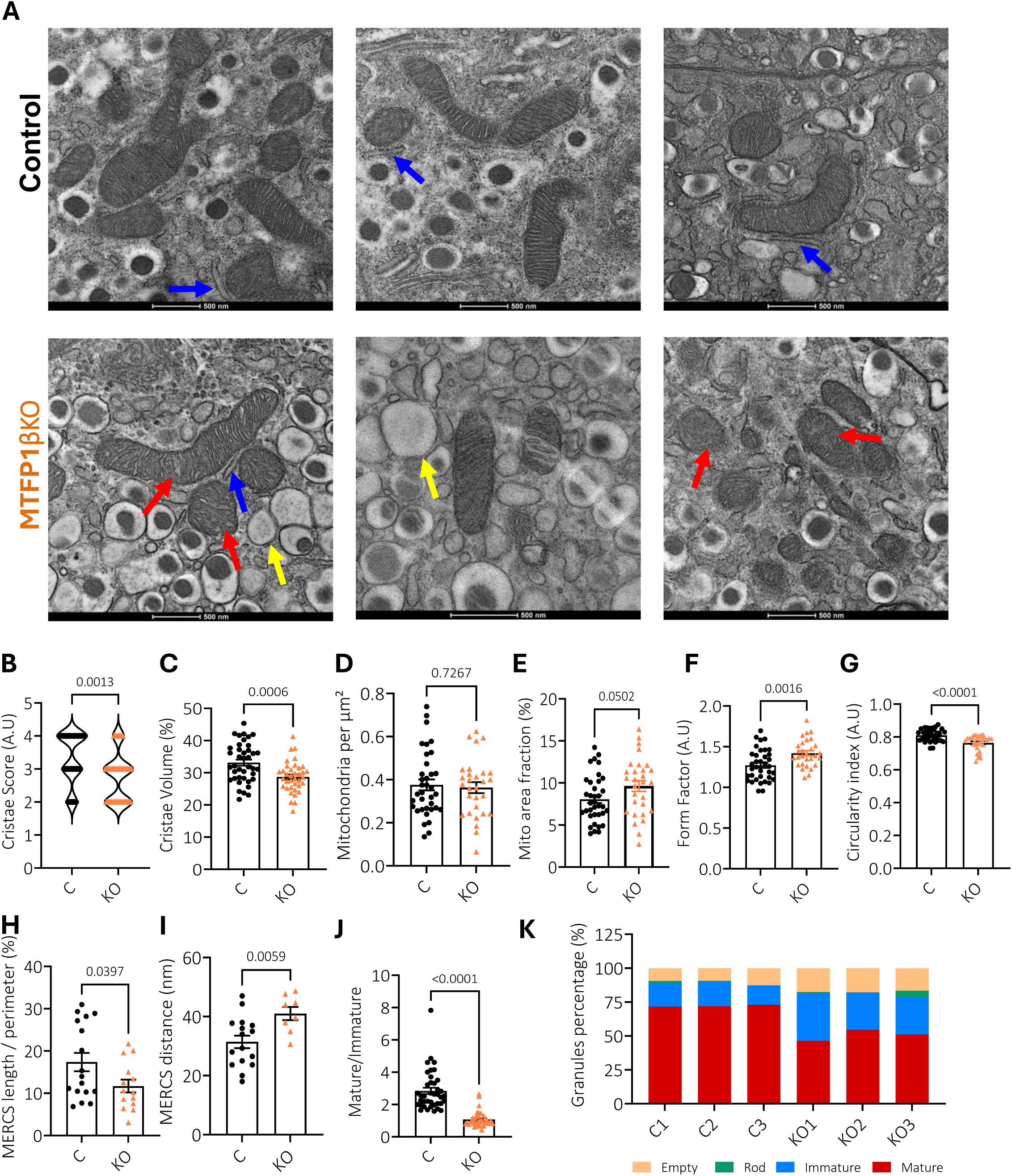
β-cell MTFP1 is essential to maintain mitochondrial cristae structure. Islets were isolated from 12-week old male control (MTFP1^f/f^) and MTFP1βKO mice (MTFP1^f/f^, InsCre^+/−^) and subjected to TEM. **A)** Representative TEM images of MTFP1βKO and control β-cells. Red arrows: distorted mitochondria with deformed cristae. Blue arrows: MERCs. Yellow arrows: non-mature/crystallized insulin granules, showing a grey or empty interior. Scale bar = 500 nm. **B-K)** Quantification of parameters: **B)** Cristae score, **C)** Cristae volume (per mitochondria), **D)** number of mitochondria per β-cell area, **E)** mitochondria area, defined as % of cellular area occupied by mitochondria, **G)** Form factor, as a measure of branching, **H)** MERCS length/ mitochondria perimeter, **I)** MERCS length of contact between the two organelles, **J)** Proportion of mature to immature granules and **K)** granule distribution within each animal. For (C), each dot represents a single mitochondrion. For all other graphs, each dot represents a single image acquisition from 3 different mice per genotype. Error bars represent SEM, Welch’s t test.

Interestingly, we also observed a significant lowering in the number of mitochondria-ER contact sites (MERCS), defined as the percentage of mitochondria membrane in contact (within 2-20 nm) with ER (Figure 6H) and the average distance between the two organelles strikingly increased (Figure 6I). Further analysis also unveiled an accumulation of empty, rod-shaped and/or uncrystallised insulin granules, with a striking reduction in the ratio of mature to immature granules in MTFP1-βKO β-cells in comparison to controls (Figure 6J, K), consistent with defective proinsulin processing

Overall, these results demonstrate that, in primary β-cells, elimination of MTFP1 leads to mitochondria that are slightly longer and more branched, with disorganised cristae that occupy a smaller proportion of the mitochondria, and reduced ER-mitochondria coupling. The combination of altered cristae architecture and increased MERCS distance likely disrupts the spatial organisation required for efficient oxidative phosphorylation and Ca²⁺ exchange between the two organelles, thereby contributing to the observed impairments in mitochondrial respiration and ATP production. Additionally, the accumulation of immature insulin granules, as well as the previously shown reduction in mature insulin content, points to broader defects in secretory maturation, possibly arising from disrupted mitochondrial organisation and bioenergetics[35].

### MTFP1 elimination triggers minimal β-cell transcriptional remodelling

Multiple reports have shown that defective mitochondrial function can result in profound changes in gene expression and loss of β-cell identity. For example, very recently, Walker et al. [28] demonstrated that in various models of mitochondrial damage—including impaired mitophagy, disrupted mitochondrial genome integrity, or defective mitochondrial fusion—OXPHOS dysfunction activates a mitochondrial integrated stress response that ultimately drives loss of β-cell identity and hyperglycaemia.

To determine whether a similar transcriptional reprogramming occurred upon loss of MTFP1, we performed bulk RNA sequencing on isolated MTFP1-βKO islets. In contrast to models in which severe mitochondrial dysfunction triggers widespread transcriptional rewiring, MTFP1 deletion resulted in remarkably limited changes in global gene expression, with only 25 differentially expressed transcripts (p adj ≤ 0.1; Figure 7A, Supplemental Table 1). Notably, several of the upregulated genes encode mitochondrial or OXPHOS-related proteins, including the nuclear-encoded mitochondrial genes *Atp5mk*, *Cox7c*, *Ndufb1*, *Timm8b*, and *Mtln*, as well as the mitochondrially encoded gene *mt-Nd3*, with only one gene with mitochondrial function, *Aldh1l2*, being downregulated. This is consistent with a compensatory transcriptional adaptation rather than a causal mechanism underlying the observed respiratory defects, and in line with our western blot analyses showing unchanged OXPHOS proteins abundance.

**Figure 7.**
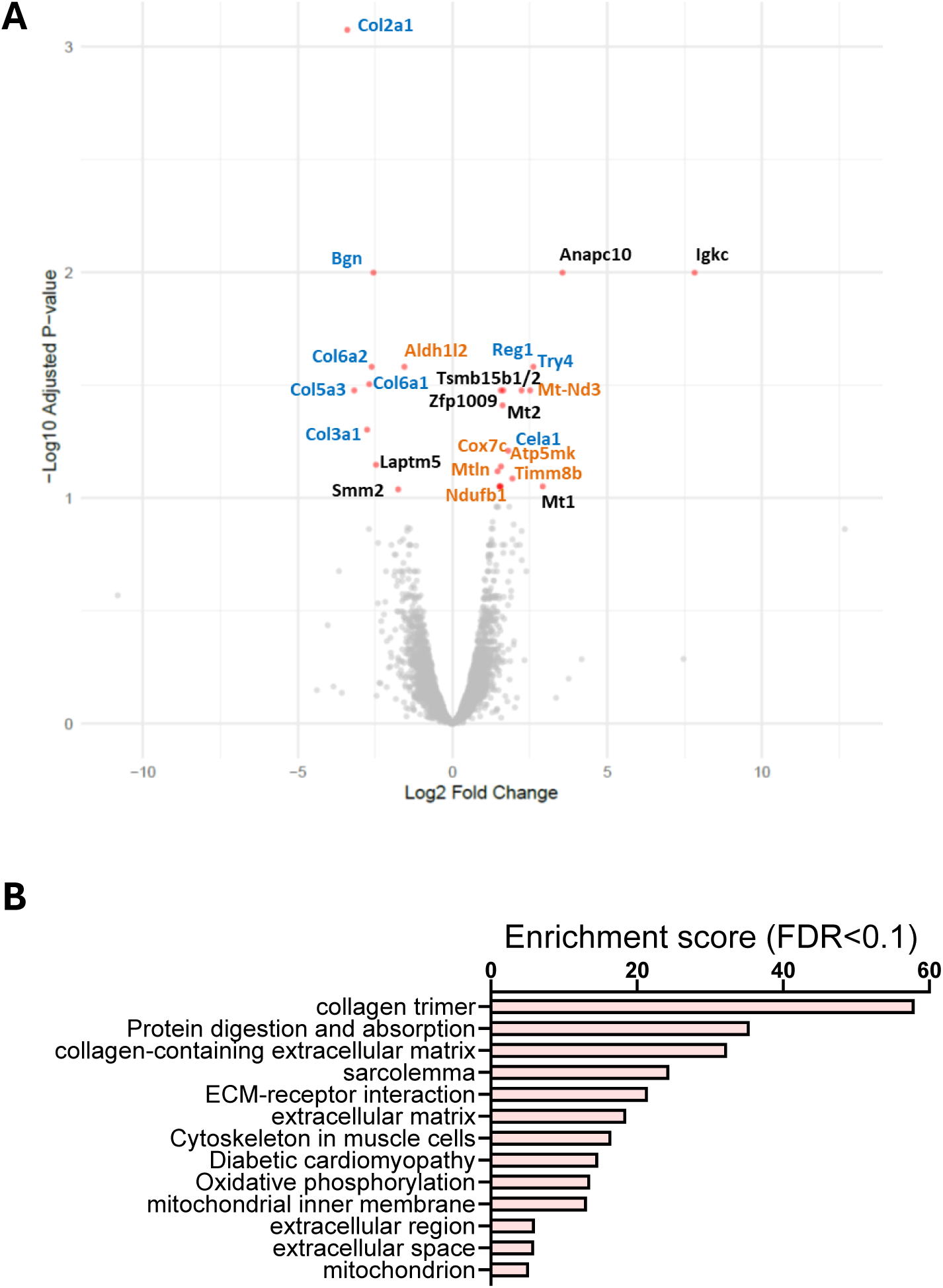
β-cell MTFP1 elimination triggers minimal transcriptional changes. Differential expression analysis was performed on RNA-seq from 12-week old male control (MTFP1^f/f^) and MTFP1βKO mice (MTFP1^f/f^, InsCre^+/−^) isolated islets. **A)** Volcano plot showing significantly (p adj < 0.1) dysregulated genes. Genes annotated to OXPHOS-related processes and extracellular matrix compartment are in orange and blue, respectively. **B)** GO analysis of significantly (P adj < 0.1) dysregulated genes performed with the Database for Annotation, Visualization and Integrated Discovery. The graph shows enrichment scores for biological processes, cellular components and KEGG’s biological pathways terms with enrichment FDR < 0.10. (See supplemental tables 1 and 2 for the full list of dysregulated genes and enriched terms).

Conversely, we observed a robust downregulation of genes encoding extracellular matrix (ECM) components and re-modellers, including *Col1a2*, *Col5a3*, *Col3a1*, *Col6a1/2*, and *Bgn*. As these transcripts are typically enriched in mesenchymal or stromal cells rather than endocrine cells, their reduction most likely reflects changes in islet microenvironment or cell–matrix interactions, rather than a loss of β-cell identity.

Gene ontology analysis further confirmed enrichment for pathways related to ECM organisation, mitochondrial inner membrane components, and oxidative phosphorylation (Figure 7B; Supplemental Table 2).

Together, these findings indicate that although MTFP1 is essential for mitochondrial architecture and bioenergetic output, its loss does not elicit the broad dedifferentiation-associated transcriptional signature characteristic of severe mitochondrial stress. Instead, MTFP1 deficiency primarily impairs β-cell function through structural disruption of the inner mitochondrial membrane and defective metabolic coupling, rather than through transcriptionally mediated loss of β-cell identity.

### Loss of MTFP1 impairs first-phase Ca²⁺ responses and reshapes β-cell connectivity

Given that MERCs are key hubs for Ca²⁺ exchange and metabolic coupling between β-cells [36, 37], we further investigated whether elimination of MTFP1 disrupted the spatiotemporal organisation of Ca²⁺ signalling and network connectivity.

Thus, we monitored glucose- and KCl-induced Ca²⁺ dynamics in intact islets incubated with the Ca²⁺ sensor Cal-520 using confocal imaging. Overall, MTFP1-βKO islets showed a striking reduction in glucose-induced changes in intracellular Ca^2+^, whilst KCl-induced increases in Ca^2+^ remained comparable (Figure 8A). This suggests that voltage-gated calcium channels remained functional and the effect of MTFP1 elimination occurred upstream of plasma membrane depolarization, possibly due to the reduced capacity of these cells to produce ATP.

**Figure 8.**
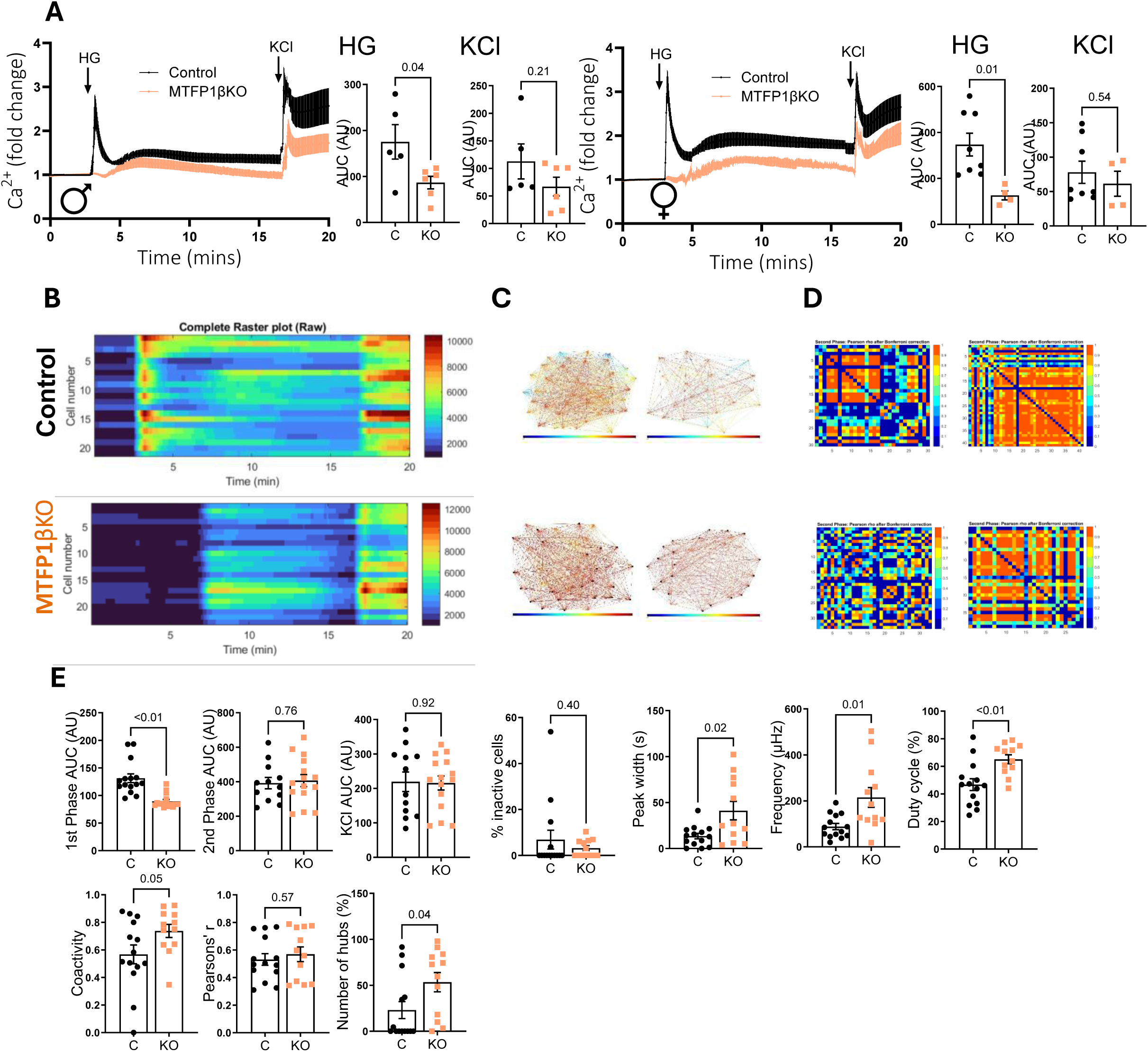
Loss of MTFP1 impairs first-phase Ca²⁺ responses and reshapes β-cell connectivity. Islets were isolated from 12-week-old male and female control and MTFP1βKO mice and subjected to real-time Ca²⁺ imaging using Cal-520 in response to high glucose (HG = 17 mM) and KCl (20 mM). **A)** Average Ca²⁺ traces (fold change over baseline), with quantification of area under the curve (AUC) for the HG and KCl responses. Each dot represents a single acquisition (n=5-6), with 3-8 islets per acquisition isolated from 3 different mice/genotype. **B-E)** Functional network analysis of β-cell activity in individual islet cells. **B)** Representative raster plots of Ca²⁺ activity (raw heatmaps) from a control and a MTFP1βKO islet. **C)** Representative coactivity plots illustrating cell-cell connectivity across single islet cells in two control and two MTFP1βKO islets. **D)** cell-cell pairwise correlation matrices depicting Bonferroni-corrected Pearson’s r for the whole Ca²⁺ traces within two control and two MTFP1βKO islet. **E)** Statistical comparison of the properties of the calcium traces measured in individual cells between the control and a MTFP1βKO islets, including first-phase and second-phase HG responses (AUC, area under the curve), KCl AUC and percentage of inactive cells, peak width, oscillation frequency and duty cycle, together with network parameters (coactivity index, Bonferroni-Pearson’s r, and number of hubs), calculated considering the second phase only. Each dot represents a single islet (n=15 islets/genotype) from 3 mice/genotype. Error bars represent SEM, unpaired Student’s or Welsch’s t test.

Detailed analysis of Ca^2+^ changes at the single-cell level in intact control islets confirmed that glucose stimulation elicited a rapid and robust increases, initiating in a subset of first-responder cells and propagating through the islet as well-synchronised oscillatory waves (Figure 8B). In contrast, MTFP1βKO islets exhibited a markedly blunted first-phase Ca²⁺ response, with delayed activation and reduced amplitude of the initial rise (Figure 8B). Despite this attenuated onset, the subsequent Ca²⁺ oscillations appeared more co-active, or connected, across cells, as reflected by denser correlation matrices and network graphs (Figure 8C, D). Quantitative analysis confirmed a significant reduction in the number of first-responder cells, alongside an unexpected increase in overall functional connectivity (Figure 8E). The strength and density of cell–cell correlations were both elevated in MTFP1βKO islets compared with controls (Figure 8E), raising the possibility that impaired MERCS and mitochondrial Ca²⁺ handling dampen early metabolic responsiveness but promote compensatory synchronisation across the β-cell network. Together, these findings indicate that MTFP1 is required for the initiation and hierarchical propagation of Ca²⁺ signals within the islet, and that its loss later promotes network organisation, leading to delayed but more uniform Ca²⁺ activity.

In summary, we have uncovered an essential role for MTFP1 in pancreatic β-cells, establishing it as a previously unrecognised regulator of mitochondrial inner membrane architecture essential for efficient ATP production, Ca²⁺ signalling and glucose-stimulated insulin secretion.

### Context-dependent and β-cell–specific regulation of MTFP1 under metabolic stress

We have previously shown that miR-125b expression in islets is increased during exposure to high levels of glucose *in vitro* via inactivation of AMP-activated kinase[25], prompting us to test whether MTFP1 expression is lowered in islets in response to high glucose. However, treatment of both mouse and human islets with increasing glucose concentrations, did not affect MTFP1 protein expression or mRNA levels (Supplemental Figure 5). This suggested additional levels of regulation and/or possibly reflects the relatively small changes occurring in miR-125b expression under these conditions[25].

To determine whether a longer-term, *in vivo*, exposure to glucolipotoxicity alters MTFP1 expression, we fed C57BL6/J mice a high-fat, high-sugar diet for 12 weeks, which, as expected, resulted in significantly increased body weight and impaired glucose tolerance (Figure 9A, B). Consistent with our earlier *in vitro* findings[25], islets from these mice displayed higher levels of miR-125b (Figure 9C) and lowered AMPK Thr172-phosphorylation, implying inhibition of the enzyme (Figure 9D). Nevertheless, and contrary to what we anticipated, MTFP1 expression was up-regulated (Figure 9D), whereas no significant changes were detected in Mtfp1 mRNA (Figure 9E). Moreover, although no correlation was observed between miR-125b expression and MTFP1 protein levels in the islets of these animals, a negative, albeit non-significant correlation was detected with AMPK phosphorylation (Figure 9F). On the other hand, a striking correlation was observed between the extent of weight gain of the animals and MTFP1 expression (r=0.7819, p=0.006, Figure 9F), suggesting that physiological changes accompanying weight gain may activate compensatory mechanisms that increase MTFP1 expression independently of miR-125b. Notably, MTFP1 translation has been reported to be stimulated by mTOR[38], which is itself negatively regulated by AMPK[39], which may favour increased MTFP1 protein abundance under conditions of reduced AMPK activity.

**Figure 9.**
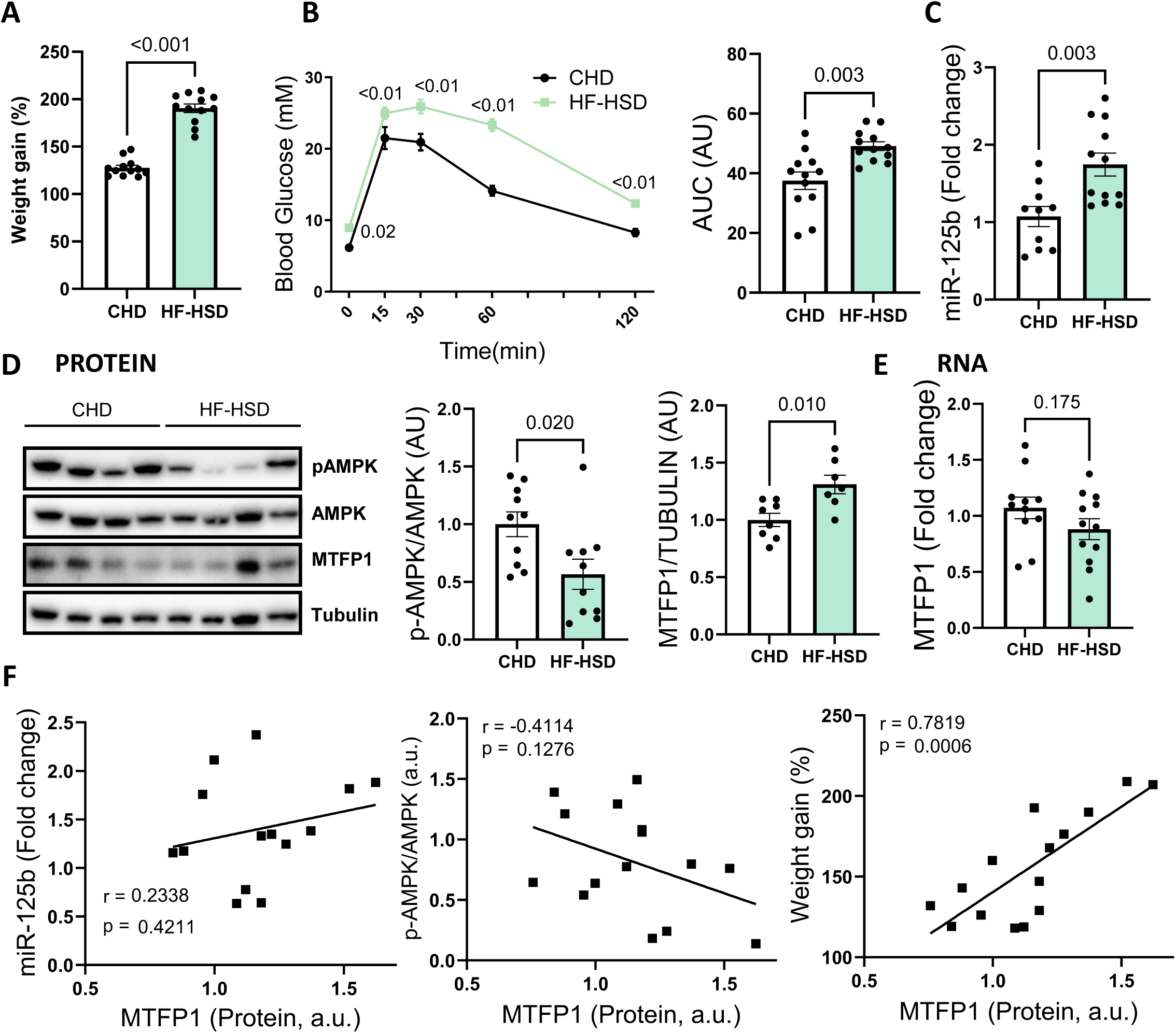
MTFP1 protein levels are increased in islets from mice fed a high fat, high sugar diet for 12 weeks and directly correlate with weigh gain. 7 week-old C57BL6/J male mice were fed a chow (CHD) or a high-fat, high-sugar (HF-HSD) diet for 12 weeks ad libitum. A) Weight gain (left-hand side panel) and B) oral glucose (2g/kg) tolerance test (middle panel, with area under the curve (AUC) in the right-hand side panel). C) MiR-125b quantitative RT-qPCR in isolated islets, normalized to that of the endogenous control miR-574-3p [26]. D) Representative Western blot showing protein levels of phospho-AMPK (Thr172, pAMPK), total AMPK (AMPK), MTFP1 and tubulin. Bar graphs show densitometry quantification of pAMPK relative to total AMPK and MTFP1 normalized by Tubulin. Data is presented relative to the average CHD value. E) *Mtfp1* mRNA quantitative RT-qPCR in isolated islets, normalized to *Ppia* mRNA and presented as relative to the average CHD value. F) MTFP1 protein expression data plotted against miR-125b expression (left-hand side panel), pAMPK (middle panel) and weight gain (right-hand side panel) in the same mouse islets and showing results from Pearson correlation analysis (r, p). Each dot represents a single mouse (n=8-14/diet); AU = arbitrary units. Error bars represent SEM, unpaired t test with Welch’s correction (A, B (AUC), C, D), two-way ANOVA (repeated measures) and Fisher’s LSD (B) and simple linear regression (F).

Additionally, we surveyed data from the human islet project, recently made available at https://www.humanislet.com ([40], Alberta Diabetes Institute IsletCore) and found a lack of correlation between *MTFP1* mRNA and protein expression in human islets and the body mass index (BMI) of the donors (data not shown). Intriguingly, a strong negative correlation was identified between pseudobulk *MTFP1* gene expression levels in β-cells and HbA1c of the donors (coefficient= -4.8, p=0.0879, Supplemental Figure 6A). Moreover, pseudobulk *MTFP1* gene expression analysis identified a possible downregulation of *MTFP1* expression in β-cells from donors with pre-diabetes (defined as HbA1c of 5.7-6.5% and no clinical diagnosis of T2D) and T2D in comparison with healthy individuals. These tendencies were not apparent in whole islet bulk gene expression data (Supplemental Figure 6A,B). Although these results will need to be verified in a higher number of samples, they suggest a β-cell-specific effect of glucose on MTFP1 expression, emphasising the need to study MTFP1 expression in isolated β-cell populations, or single cells, rather than in full islets in future studies.

## Discussion

In this study, we identify MTFP1 as a central regulator of β-cell mitochondrial architecture, bioenergetics and insulin secretory capacity. By combining genetic loss- and gain-of-function approaches with high-resolution imaging and detailed physiological analyses, we demonstrate that β-cell MTFP1 is required to maintain cristae integrity, OXPHOS efficiency, and ER–mitochondria coupling, thereby enabling efficient ATP production, Ca²⁺ responses, and glucose-stimulated insulin secretion.

### MTFP1 role beyond fission/fusion in β-cells

Our data support a model in which MTFP1 acts primarily as a regulator of inner mitochondrial membrane (IMM) organisation and proton motive force in β-cells. Across complementary *in vitro*, *ex vivo* and *in vivo* models, loss of MTFP1 produced only modest and context-dependent changes in mitochondrial length and circularity, yet consistently resulted in profound defects in mitochondrial respiration and GSIS. In β-cell–specific MTFP1 knockout islets, mitochondrial elongation was only mildly affected, yet basal and maximal respiration, ATP-linked OCR, membrane potential, ATP generation and first-phase Ca²⁺ responses were all markedly impaired. Together, these findings indicate that in β-cells, MTFP1 is critical for maintaining IMM structure, cristae organisation and mitochondria–ER contact sites (MERCs).

This conclusion is consistent with and extends recent *in vivo* work on MTFP1 in other tissues. In cardiomyocytes, Donnarumma et al. [22] showed that deletion of *Mtfp1* causes adult-onset dilated cardiomyopathy associated with specific IMM defects, including futile proton leak dependent on adenine nucleotide translocase and increased sensitivity to mitochondrial permeability transition pore (mPTP) opening, without major changes in mitochondrial division, indicating that MTFP1 is dispensable for fission but essential for IMM integrity and bioenergetic efficiency. Altered cristae structure was also reported in symptomatic, but not pre-symptomatic, MTFP1 knockout hearts, raising the possibility of secondary of cardiomyopathy. In our study, MTFP1-dependent cristae and bioenergetic defects were detected in β-cells from glucose-intolerant, non-hyperglycaemic mice, arguing against hyperglycaemia-driven secondary mitochondrial damage. By contrast to the deleterious effects observed in heart and β-cells, hepatocyte-specific *Mtfp1* ablation enhances hepatic OXPHOS and mitochondrial respiration and protects against high-fat-diet–induced steatosis and systemic metabolic dysregulation, again in the absence of overt morphological changes, nor cristae disorganisation[41]. Together with our data, these studies support the view that MTFP1 is a context-dependent tuner of IMM properties whose mechanistic and functional output (its elimination is deleterious in heart and β-cells, and protective in liver) depends on cell-specific metabolic demands and vulnerability to changes in proton motive force and mitochondrial membrane potential. This study further highlights the cell-type specific effects of MTFP1.

Besides the modest effects on mitochondrial elongation, there are further contrasts between our findings and the recent work by Tabara et al.[24] suggesting that MTFP1 negatively regulates IMM fusion and mitophagy in U2OS cells, where its loss permits fusion of damaged mitochondria and increases mtDNA content. In our β-cell models, MTFP1 modulation did not measurably alter mtDNA copy number, despite clear effects on respiration. This suggests that the balance between IMM fusion and mtDNA maintenance may differ substantially between β-cells and U2OS cells, reinforcing the view that MTFP1 function is strongly cell-type specific and shaped by metabolic demand.

Further supporting a major role for MTFP1 in regulating β-cell OXPHOS independently of classical mitochondrial dynamics is the observation that β-cell–specific deletion of Mfn1/2—rather than producing an opposite phenotype—similarly leads to impaired mitochondrial function, reduced mitochondrial membrane potential, and severely blunted GSIS, despite involving marked mtDNA depletion and pronounced mitochondrial fragmentation/fusion defects[29, 42]. Similarly, selective elimination of *Drp1* in pancreatic β-cells led to impaired insulin secretion and marked abnormal mitochondrial morphology, without any observable defects in oxygen consumption rates[13].

Another notable effect of MTFP1 elimination was the pronounced reduction in mitochondrial membrane potential, which could lead to mitophagy via PINK1/Parkin sensing[43]. However, such an activation would typically be accompanied by a lowering of mitochondrial number or area, which was not observed in any of our β-cell models of MTFP1 loss-of-function, including MTFP1βKO islets. One untested possibility is that loss of MTFP1 may simultaneously impair mitophagy, as MTFP1 has previously been proposed to function as a mitophagy receptor in certain cell types, including oral carcinoma Cal33 and osteosarcoma U2OS cells, through interaction with LC3-II[24, 44]. It is also possible that the accumulation of dysfunctional mitochondria overwhelmed the mitophagic capacity of the MTFP1-deficient cells, as reported in islets exposed to HFD-induced metabolic stress, where mitophagy is strongly induced yet mitochondrial dysfunction persists [45].

It is also conceivable that a more striking role for MTFP1 in regulating mitochondrial dynamics would emerge under certain physiological states that favour mitochondrial fusion in β-cells. However, such conditions remain poorly defined, as maximal mitochondrial network fusion seems to occur at physiological glucose concentrations, whereas both chronic starvation and glucolipotoxic stress favour fragmented networks[46, 47]. In addition, our imaging approach relied on diffraction-limited confocal microscopy rather than super-resolution techniques; thus, rapid or small-scale IMM remodelling events—particularly those occurring at cristae junctions or MERC interfaces—may have gone undetected. It therefore remains possible that MTFP1 influences dynamic mitochondrial rearrangements in β-cells that are too transient or below the spatial resolution of our imaging modalities to manifest as major morphological changes at steady state.

### MTFP1 loss induces energetic uncoupling without β-cell dedifferentiation – implications for β-cell failure in diabetes

Mitochondrial dysfunction is widely recognised as a central driver of β-cell failure in T2D, with numerous studies documenting defects in OXPHOS, ATP generation and mitochondrial morphology in human T2D islets and in rodent models of diabetes (reviewed in [1]). Anello et al. [4] reported that β-cells from T2D subjects exhibit impaired GSIS associated with reduced ATP levels, decreased ATP/ADP ratio, impaired mitochondrial hyperpolarisation and characteristic alterations in mitochondrial structure, including increased volume and abnormal cristae. More recent imaging and ultrastructural analyses in humans have reinforced the view that altered cristae morphology is tightly associated with reduced GSIS and β-cell dysfunction[1, 42].

Our findings add MTFP1 to the growing list of mitochondrial proteins that link structural changes in β-cell mitochondria to functional defects in insulin secretion. Similar to the situation in T2D islets, MTFP1-deficient β-cells show impaired ATP production, reduced membrane potential and altered cristae architecture, together with diminished insulin secretion and Ca²⁺ responses. But in contrast to many models of β-cell mitochondrial dysfunction, including those involving impaired mitophagy, disrupted mitochondrial genome integrity, or deletion of fusion proteins such as *Clec16a*, *Tfam* and *Mnf1/2* [28, 29], impaired OXPHOS did not trigger a broad transcriptomic rewiring and loss of β-cell identity. Indeed, only 25 transcripts were significantly altered, including six genes encoding proteins annotated to mitochondrial/OXPHOS pathways (consistent with a compensatory adaptation), as well as a subset of downregulated extracellular matrix genes, pointing to microenvironmental rather than β-cell identity changes. Notably, a similar dissociation between pronounced mitochondrial functional defects and limited transcriptional remodelling was recently reported following liver-specific deletion of MTFP1 under basal, show-fed conditions[41]. This resembles more a “structural uncoupling” of mitochondria from their energetic and signalling context than a full mitochondrial integrated stress response (mt-ISR)-driven dedifferentiation, as observed in the aforementioned models, and suggests that not all forms of OXPHOS impairment in β-cells converge on the same transcriptional outcome.

The most plausible explanation for these differences is the absence of detectable effects in mtDNA content following MTFP gain- or loss-of-function in β-cells. This distinction is important when positioning MTFP1 within the broader landscape of mitochondrial defects linked to T2D: MTFP1 deficiency may represent a distinct class of β-cell mitochondrial dysfunction in which cells remain alive, transcriptionally “β-like”, yet metabolically uncoupled from glucose due to defective cristae architecture and impaired ATP generation.

In this context, the upregulation of MTFP1 protein observed in islets from mice exposed to high-fat, high-sugar diet, and its strong correlation with weight gain, likely reflects an early compensatory attempt in β-cells to meet rising insulin demand. This is fully consistent with our functional data, where enforced MTFP1 expression in β-cell lines and primary human islets enhances GSIS, indicating that β-cells gain secretory efficiency when MTFP1 levels rise. Thus, mouse β-cells appear capable of elevating MTFP1 expression as a protective response to increased metabolic load, helping maintain ATP production and secretory competence. By contrast, the pseudobulk β-cell transcriptomic data from human donors suggest a different, potentially later stage of β-cell damage, in which MTFP1 mRNA tends to decline in β-cells from pre-diabetic and T2D individuals. Rather than contradicting the compensatory pattern observed in mice, this may reflect the point at which β-cell compensatory mechanisms begin to fail, possibly due to chronic glucolipotoxicity, inflammatory stress, or dedifferentiation cues that limit the capacity to upregulate MTFP1. However, this trend in human β-cells is based on modest sample numbers and should be interpreted with caution.

Although, as noted above, most (6 out of 7) dysregulated genes annotated to mitochondrial inner membrane processes such as oxidative phosphorylation were upregulated, consistent with compensatory adaptation to mitochondrial dysfunction, *Aldh1l2*, encoding for mitochondrial 10-formyltetrahydrofolate dehydrogenase, was strongly downregulated (log2 Fold change= 1.6, padj= 0.03). ALDH1L2 is an important source of folate-dependent NADPH within mitochondria[48]; however, the functional significance of its regulation in β-cells remains unclear, as the role of this enzyme has not, to our knowledge, been studied in this cellular context. Notably, treatment of islets with the secregatogue dextrorphan (DXO) has been shown to induce *Aldh1l2* expression via activating transcription factor 4 (ATF4), resulting in an increased mitochondrial NADPH/NADP+ ratio through activation of serine-linked mitochondrial one-carbon metabolism. This response was proposed to support islet cell protection during oxidative stress, albeit at the expense of insulin secretory function[49].

The enrichment of downregulated ECM genes was unexpected and is likely to reflect the presence of non-islet cells, such as stellate and endothelial cells, within isolated islets[50, 51]. The ECM, including that of the basement membrane and interstitial matrix, is a key component of the islet microenvironment that is essential for maintaining normal islet morphology and function and is altered in both T2D and T1D [52]. It is conceptually possible that β-cell autonomous defects induced by MTFP1 loss exert secondary effects in the islet microenvironment, as β-cell secretory activity, including the release of both insulin and ATP, has been shown to influence the proliferation and signalling of stellate, vascular and immune cells within and surrounding the islets, as elegantly summarised in [52]. Nevertheless, as the process of islet isolation using collagenase is likely to disrupt ECM integrity and associated cell populations, further investigating this possibility will require *in* vivo approaches and is beyond the scope of the present study.

### MTFP1, cristae, MERCs and β-cell network behaviour

An unexpected finding of this work was the striking reduction in ER–mitochondria contact length, together with an increased inter-organelle distance. This finding is also supportive of a different mechanism of action than that of classical regulators of β-cell mitochondrial fission and fusion. For example, *Mfn1/2*-defficient β-cells preserved mitochondrial-ER proximity[29], though their ablation has been shown to alter MERCs in other cellular contexts[53].

Mitochondrial cristae shape and density are well-established determinants of respiratory chain supercomplex assembly and ATP synthase efficiency, and even subtle alterations in cristae organisation can exert disproportionately large effects on local ΔΨm and ATP production [54, 55]. Moreover, although the outer mitochondrial membrane is permeable to ions, the inner membrane is not; thus, the architecture of cristae and their junctions directly influence how protons and Ca²⁺ accumulate within the mitochondrial matrix. In this context, the Mitochondrial Contact Site and Cristae Organizing System (MICOS) complex has emerged as a central regulator of cristae junction formation and inner–outer membrane contact sites, and therefore as an important modulator of IMM-dependent signalling processes—including mitochondrial Ca²⁺ handling—complementing the more established role of the mitochondrial Ca²⁺ uniporter complex [55–57]. Notably, several components of the MICOS complex have been identified among potential interactors of MTFP1 in U2OS cells[24]. It is therefore possible that defective Ca²⁺ uptake by β-cell mitochondria further impairs their capacity to generate ATP. Moreover, MERCs are now recognised as key hubs for bidirectional Ca²⁺ flux, phospholipid exchange and metabolic signalling, and have been implicated in systemic glucose homeostasis and diabetes pathogenesis[36, 37]. Together, the structural changes we report provide a plausible explanation for the concurrent reductions in membrane potential, ATP-linked respiration and glucose-induced ATP dynamics that we observe, as well as for the blunted first-phase Ca²⁺ response.

Paradoxically, despite displaying a markedly delayed and blunted initial Ca²⁺ rise, MTFP1-βKO islets showed a second-phase response characterised by increased overall functional connectivity and wider and more frequent Ca²⁺ oscillations. Given the emerging role of ER Ca²⁺ in supporting β-cell synchrony [58], an interesting possibility is that the MERCs reduction lowers ER to mitochondria Ca²⁺ transfer, which instead further contributes to cytosolic Ca²⁺ oscillations. Moreover, reduced ER-Ca²⁺ into mitochondria transfer could also contribute to preservation of β-cell mass despite secretory dysfunction, based on previous observations that proteins important for MERCs formation such as PDZ domain-containing 8 (PDZD8) promote β-cell death in diabetes by enhancing ER-mitochondria contacts and facilitating Ca²⁺ transfer[59].

A reduction in ER–mitochondrial interactions has been reported in β-cells from individuals with type 2 diabetes (T2D)[60]. More recently, these contact sites have been shown to act as hotspots for interaction with endosomal GLP-1 receptors following functional stimulation with GLP-1R agonists, therapies widely used in the treatment of T2D, thereby triggering changes in mitochondrial function and remodelling[10]. Whether MTFP1 plays a role in these processes, or whether reinforcing MTFP1 expression could restore these responses, remains to be determined.

### A major role for MTFP1 as a downstream effector of miR-125b

An important outcome of this work is the identification of *MTFP1* as a major downstream effector of miR-125b in β-cells. We previously showed that miR-125b is induced by hyperglycaemia via AMPK and acts as a negative regulator of β-cell function, whereas its reduction in human islets and a β-cell line improved their secretory capacity[25]. It is important to note that the phenotype we observed upon β-cell selective miR-125b overexpression[25] is not fully mirrored by the MTFP1-βKO animals generated here. For instance, MTFP1-βKO showed a more delayed, milder glucose intolerance compared to MIR125B-Tg mice. Additionally, MIR125B-Tg islets displayed striking transcriptional changes, in sharp contrast with the little effect observed in MTFP1-βKO islets. This is not surprising given the pleiotropic roles of miRNAs and multiple targeting[61], with our high-throughput experimental pipeline identifying dozens of potential miR-125b targets in MIN6 insulinoma cells[25]. Nevertheless, the two models share some important features, including defective maturation of insulin granules and lower levels of mature insulin. Whilst other miR-125b targets such as the mannose-6-phosphate receptor may contribute to these insulin maturation defects in MIR125B-Tg mice through the regulation of lysosomal function and membrane trafficking, as previously discussed[25], it is also possible that MTFP1-mediated impairment of mitochondrial function plays a major role. For example, lowered mitochondrial ATP production may impair insulin granule maturation by limiting ATP-dependent V-ATPase–driven granule acidification and thus compromising the optimal pH/Ca^2+^ milieu for prohormone convertases[62] or by perturbing ATP-dependent ER proteostasis and Ca^2+^ or ER redox homeostasis upstream of granule formation[35, 63]. Additionally, in EndoC-βH1 cells, the enhanced respiration, mitochondrial shortening and improved GSIS observed upon miR-125b deletion are fully dependent on MTFP1 upregulation, and MTFP1 knockdown reverses these beneficial effects. This places MTFP1 downstream of a glucose–AMPK–miR-125b axis that modulates β-cell mitochondrial fitness. Nevertheless, in islets from mice exposed long-term to a high-fat, high-sugar diet, we find increased miR-125b levels and reduced AMPK phosphorylation, as expected, but MTFP1 protein expression is paradoxically increased without changes in Mtfp1 mRNA. This suggests that, *in vivo*, MTFP1 is regulated by additional post-transcriptional or post-translational mechanisms that can offset miR-125b-mediated repression under metabolic stress, potentially as part of a compensatory response to the increased load placed on β-cell mitochondria and the ER. This idea is supported by the strong correlation between weight gain and MTFP1 levels and by our MTFP1 overexpression experiments leading to improved secretory capacity of human β-cell lines and islets. The divergent associations between MTFP1 expression and HbA1c in bulk vs β-cell–specific human islet datasets further support the idea that MTFP1 regulation is both cell-type specific and stage dependent. Future work will be required to dissect these regulatory layers and to determine whether MTFP1 upregulation in pre-diabetic and T2D β-cells is adaptive, maladaptive, or both. If the former holds true, preventing miR-125b binding to MTFP1 mRNA may represent a novel therapeutic strategy to enhance MTFP1 expression and, consequently, β-cell function in T2D.

Overall, our work establishes MTFP1 as a previously unrecognised, β-cell–specific regulator of IMM architecture, MERCs and bioenergetic output, required for first-phase Ca²⁺ responses, coordinated β-cell network activity and effective insulin secretion. In contrast to other forms of mitochondrial dysfunction that drive β-cell dedifferentiation via mt-ISR, MTFP1 loss primarily induces a structural and functional uncoupling of mitochondria from their metabolic and signalling milieu, with limited effects on β-cell identity. When integrated with studies in heart and liver, MTFP1 emerges as a context-dependent regulator whose precise role is dictated by tissue-specific demands on oxidative metabolism and Ca²⁺ handling. Future work should address how MTFP1 is regulated across the spectrum from health to T2D in human β-cells, whether targeted modulation of MTFP1 or its upstream regulators (including miR-125b) can restore mitochondrial fitness in dysfunctional islets, and how MTFP1 interfaces with other mitochondrial stress pathways and contact site regulators that have been implicated in diabetes.

## Materials and Methods

### Cells and islets culture

HEK-293T and AD293 cells were cultured in high-glucose (4.5 g/L) DMEM (Gibco) supplemented with 10% FBS, 2 mM L-glutamine and 100 U/mL penicillin–streptomycin (P/S). EndoC-βH3 cells[27] were cultured in Advanced DMEM/F12 containing 2% BSA Fraction V, 50 μM 2-mercaptoethanol, 2 mM L-glutamine, 10 mM nicotinamide, 5.5 μg/mL transferrin, 6.7 ng/mL sodium selenite and 100 U/mL P/S. Plates were coated with 804G matrix[64]. To induce Cre-ERT2–mediated recombination in EndoC-βH3 cells, cultures were treated with 1 µM 4-hydroxytamoxifen (4-OHT; “tamoxifen”) for 21–42 days before functional experiments[27]. EndoC-βH1 cells[65] were cultured in low-glucose DMEM (1 g/L) supplemented with 2% BSA Fraction V (fatty acid free), 10 mM nicotinamide, 5.5 μg/mL transferrin, 6.7 ng/mL sodium selenite, 50 μM 2-mercaptoethanol and 100 U/mL P/S, on plates coated with 2 μg/mL fibronectin and 1% extracellular matrix. Mouse pancreatic islets were isolated with collagenase as described previously[66] and maintained in RPMI 1640, 10% FBS, L-glutamine and 11mM glucose. Human islets were cultured in RPMI 1640, 10% FBS, L-glutamine, 5.5 mM glucose and 0.25 μg/μL amphotericin B. Human islets were obtained through the Integrated Islet Distribution Programme (IIDP) with the characteristics of the human donors and isolation centres shown in Supplemental Table 3. All procedures conformed to the Declaration of Helsinki and to local ethical and consent regulations of the isolation centres; use of de-identified islets for research was approved by local research ethics committees.

Cells and islets were maintained at 37°C with 95% O2/ 5% CO2.

Correlations between MTFP1 expression, BMI and HbA1c were performed with publicly available human islet bulk and single-cell RNA-seq datasets from the Human Islet Project (https://www.humanislet.com; Alberta Diabetes Institute IsletCore[67]). β-cell expression estimates were correlated with clinical parameters using linear regression.

### Mouse models

Mice were housed in a pathogen–free facility at Imperial College London, under a 12-h light/dark cycle with ad libitum access to food and water. All procedures were carried out in accordance with the UK Home Office Animal Scientific Procedures Act, 1986 (Licence PP7151519). Mice were fed a regular chow diet or, when indicated a high-fat, high-sucrose diet (35.4% carbohydrate, 23% protein, and 35.85% fat, Research Diet (D12331)).

To generate β-cell–specific MTFP1 knockout mice (MTFP1-βKO), Mtfp1^lox/lox^ mice generated on a C57BL/6N background[22] were crossed with Ins1-Cre mice on a C57BL/6J background [30]. Lines were backcrossed for at least five generations to achieve >95% C57BL/6J genetic background and littermates lacking Cre (Mtfp1^lox/lox^) were used as controls.

### DNA and RNA extraction and quantitative PCR

Genomic (g) and mitochondrial (mt) DNA for mtDNA copy number was extracted from cells and islets by overnight lysis in SNET buffer (20 mM Tris-HCl pH 8.0, 5 mM EDTA, 400 mM NaCl, 1% SDS, 40 μg/mL Proteinase K) at 55°C, followed by phenol:chloroform:isoamyl alcohol extraction, isopropanol precipitation and 70% ethanol wash. For mtDNA copy number analyses, qPCR was performed using 12 ng (cells) or 8 ng (mouse islets) total DNA with SYBR Green Master Mix (Applied Biosystems) and primers targeting mitochondrial tRNA-Leu (UUR) (human), mt-ND1 (mouse) and nuclear β2-microglobulin (human) or hexokinase (mouse) (Supplemental Table 4). Relative mtDNA/gDNA content was calculated using the ΔCt method.

Total RNA from EndoC-βH1/βH3 cells and mouse and human islets was extracted using TRIzol (Invitrogen) according to the manufacturer’s instructions. For mRNA quantification, 100-500 ng RNA was reverse-transcribed using the High-Capacity cDNA Reverse Transcription Kit (Thermo Fisher). qPCR was performed on a 7500 Fast Real-Time PCR System (Applied Biosystems) using SYBR Green Master Mix and gene-specific primers (Supplemental Table 4). *MTFP1* mRNA was normalised to *RPLP0* (human) or *PPIA* (mouse) using the ΔCt method. For miRNA analysis, 30 ng RNA was reverse transcribed using the miRCURY LNA Universal RT miRNA Kit (Qiagen). qPCR was performed with miRCURY LNA SYBR Green PCR Kit and LNA primers for hsa-miR-125b-5p and reference miRNAs hsa-let-7d-3p[25], and relative expression was calculated using ΔCt.

### Generation of MTFP1 gain- and loss-of-function cellular models

EndoC-βH1 MIR125B2-KO cells were generated using paired gRNAs targeting *MIR125B2* loci as described[25]. For generation of EndoCβH3-MTFP1-KO cells, two different gRNA pairs (KO1 and KO2, Supplemental Table 5) targeting exons 1-2 of the human *MTFP1* locus were subcloned as previously described under an H1 and U6 promoter within the same lentiviral vector[25], following the substitution of the puromycin resistance gene by a blasticidine. In this vector, hSpCas9 expression is driven by a RIP (rat Insulin promoter). EndoCβ-H3 cells were infected with lentivirus expressing these gRNAs and the hSpCas9 (driven by a rat insulin promoter) and integrating cells were selected by subculture in the presence of 8 μg/ml blasticidine. Cells expressing non-targeting gRNAs were used as controls (Supplemental Table 5). RT-qPCR for *MTFP1* using primers outside the targeted region (Supplemental Table 4) was performed to assess the levels of *MTFP1* in the resultant cell populations. All the experiments were performed with controls and KO cell populations generated from at least three independent infections.

Adenovirus expressing HA-tagged, codon-optimised human MTFP1[22] (Ad-MTFP1) was generated using the AdEasy system[68] by subcloning into pAdTrack-CMV. Viral particles produced and amplified in AD293 cells and amplified. Viral particles were purified by CsCl gradient ultracentrifugation and dialysed into storage buffer (10% glycerol, 10 mM Tris-HCl, 10 mM MgCl₂, 15 mM NaCl) before storage at −80°C. Viral titres (PFU/µL) were estimated by GFP-positive plaque formation in AD293 cells. EndoC-βH3 cells were infected with Ad-MTFP1 or control GFP-only adenovirus at MOI 2 for 48 h unless otherwise indicated. Intact human islets were pretreated briefly with Accutase to improve viral penetration[69] and infected with Ad-MTFP1 or control virus at MOI 5 for 72 h.

ShRNAs targeting human MTFP1 (siRNA sequence: GCCCTTCCCTGACCCAATAAA) or non-targeting (siRNA sequence: CAACAAGATGAAGAGCACCAA) were cloned into pLKO.1 under a U6 promoter and lentivirus produced in HEK293T cells by co-transfection with psPAX2/pMD2.G, following the pLKO.1-TRC Cloning vector protocol from Addgene (https://www.addgene.org/protocols/plko/). Human islets were exposed to lentiviral particles encoding MTFP1 shRNA or non-targeting shRNA at MOI 20 for 72 h following a brief Accutase treatment as above. EndoC-βH1 control and MIR125B2-KO cells were infected at MOI 5. Knockdown was confirmed by RT-qPCR for MTFP1.

### Glucose-stimulated insulin secretion (GSIS) and insulin content

EndoC-βH1 and EndoC-βH3 cells were seeded in 48-well plates (6 × 10⁴ and 7 × 10⁴ cells/well, respectively) and infected at the indicated MOI 48h prior the experiments. Cells were pre-incubated overnight in low-glucose medium (2.8 mM). The following day, cells were equilibrated for 1 h in Krebs–Ringer–bicarbonate HEPES (KRBH: NaCl 115 mM, KCl 5 mM, NaHCO₃ 24 mM, MgCl₂ 1 mM, CaCl₂ 2.5 mM, HEPES 10 mM, 0.2% BSA, pH 7.4) containing 0.5 mM glucose at 37°C. Cells were then incubated for 1 h in KRB with 0.5 mM glucose or 15 mM glucose. Supernatants were collected and clarified by centrifugation (3000 rpm, 5 min, 4°C). Cells were lysed in TETG buffer (20 mM Tris-HCl pH 8.0, 137 mM NaCl, 1% Triton X-100, 10% glycerol, 2 mM EGTA plus protease inhibitors) to measure total insulin content.

Size-matched mouse or human islets (15 per condition, in triplicates) were pre-incubated for 1 h in KRBH containing 3 mM glucose at 37°C. Islets were then sequentially incubated for 1 h at 3 mM glucose, 17 mM glucose, and 3 mM glucose + 20 mM KCl. After each step, supernatants were collected for insulin measurement and islets were finally lysed in acidified ethanol (1.5% HCl, 75% ethanol, 0.1% Triton X-100) for insulin content.

Secreted and cellular insulin were quantified using a homogeneous time-resolved fluorescence (HTRF) insulin Ultra-Sensitive Detection Kit (Revvity) following the manufacturer’s instructions.

### *In vivo* metabolic tests

For intraperitoneal glucose tolerance tests (IPGTT), mice were fasted for ∼15 h and injected intraperitoneally with 2 g/kg D-glucose. Blood glucose was measured from the tail vein at 0, 15, 30, 60, 90 and 120 min using a handheld glucometer.

Insulin tolerance tests (ITT) were performed after 5 h fast by intraperitoneal injection of 0.7 U/kg insulin (NovoRapid) and glucose measurement at 0, 15, 30, 60, 90 and 120 min.

For in vivo GSIS, overnight-fasted mice received 3 g/kg glucose intraperitoneally and blood was collected from the tail vein into EDTA-coated tubes at 0, 5 and 15 min. Plasma was separated by centrifugation (2000 × g, 20 min, 4°C) and insulin was quantified by ELISA (Crystal Chem) as per the manufacturer’s protocol.

### Western (immuno-) blotting

Cells and islets were lysed in RIPA buffer (Thermo Scientific) supplemented with EDTA-free protease inhibitors and phosphatase inhibitors (Sigma). Protein concentration was determined by BCA assay (Thermo Scientific). Equal amounts of protein (35–50 μg; 1-10 μg for insulin) were resolved on 12% Bis-Tris gels (NuPAGE) or 16% Tricine gels (for insulin) and transferred to PVDF membranes using Towbin buffer by wet or semi-dry transfer. Membranes were blocked in 5% milk in PBS–0.1% Tween-20 (PBST) and incubated overnight at 4°C with primary antibodies against MTFP1, OXPHOS complexes I–V, AMPK and phospho-AMPK, insulin, HA tag and loading controls (vinculin or β-tubulin) (Supplemental Table 6). After washing, membranes were incubated with HRP-conjugated secondary antibodies (1:7000) for 1 h at room temperature and developed using ECL reagent (BioRad). Membranes were imaged with a ChemiDoc system and quantified by densitometry.

### 3-(4,5-dimethylthiazol-2-yl)-2,5-diphenyltetrazolium bromide (MTT) cell viability assay

EndoC-βH3 cells were seeded at 6 × 10⁴ cells/well in 96-well plates and treated as indicated. Cell viability was assessed by MTT assay (Roche). After removal of culture medium, cells were incubated with 100 µL KRB and 10 µL of 12 mM MTT solution for 4 h at 37°C. Supernatant was removed, 50 µL DMSO was added to solubilise formazan and absorbance at 540 nm was read in a plate reader.

### Mitochondrial morphology analysis

For mitochondrial morphology, EndoC-βH1/βH3 cells were plated on ECM-coated glass coverslips (24 mm) at 3 × 10⁵ cells per coverslip and infected as described. After 48 h, cells were incubated in KRBH containing and 70 nM MitoTracker Green or MitoTracker Red (Thermo Fisher) for 45 min at 37°C. Images were acquired on a spinning-disk Nikon ECLIPSE Ti microscope with a 60× oil objective using 488- or 561-nm excitation (300 ms exposure). For human islets 48 h after infection, intact islets were incubated in KRBH and 100 nM MitoTracker for 30 min, washed and plated in glass-bottom chambers. Islets were imaged on a Zeiss LSM780 confocal microscope using a 63× oil objective. Z-stacks (10–15 planes at 0.48 μm spacing) were acquired per islet.

Images were analysed in ImageJ using an in-house macro as previously described, based on Wiemerslage et al. [70] to quantify mitochondria morphological characteristics including number, area, elongation, circularity and perimeter. Quantifications were performed in z-stacks of 10 images for EndoC-βH1/3 cells and in three stacks of three images each at the top, middle and bottom of the primary islets.

### Seahorse extracellular flux analysis

Mitochondrial respiration was measured using an XFe96 Seahorse analyser (Agilent). For cell experiments, XF96 plates were coated with ECM for ≥1 h and EndoC-βH1/βH3 cells were seeded at 3–4 × 10⁴ cells/well and cultured at low-glucose medium (2.8 mM) overnight. On the day of assay, cells were washed and incubated for 1 h in Seahorse assay medium (pH 7.4) containing 1 mM pyruvate, 2 mM glutamine and 0.5 mM glucose. For islets, XF96 plates were coated with poly-D-lysine and, on the day of assay, 1.5 µL Matrigel was placed in each well. Eight to ten islets were placed onto the Matrigel drop and allowed to settle before addition of 80 µL assay medium (3 mM glucose).

For mitochondrial stress tests, ports were loaded so cells were exposed to 15 mM glucose (Port A), oligomycin (2 µM for cells, 5 µM for islets; Port B), FCCP (2 µM; Port C) and rotenone/antimycin A (2.5 µM; Port D). Basal oxygen consumption rate (OCR) was measured at 3mM glucose, followed by sequential injections, with 3–4 measurement cycles per step.

After the assay, cells were fixed in 4% paraformaldehyde and stained with DAPI; total nuclear fluorescence, quantified using a Pherastar plate reader, per well was used to normalise OCR. For islets, bright-field images were acquired post-fixation, islet radius and volume were estimated assuming spherical geometry, and OCR was normalised to islet volume.

### Intracellular Ca²⁺ and ATP imaging

Cytosolic ATP/ADP dynamics were monitored in intact islets infected with adenovirus encoding PercevalHR 48h post-infection, and intracellular Ca²⁺ dynamics were assessed in intact islets loaded with Cal-520 AM (4.5μm, Stratech) as previously described[25]. Islets were incubated in KRBH for 45 min, then placed in glass-bottom chambers in BSA-free KRBH. Time-lapse imaging was performed on a Nikon ECLIPSE Ti spinning-disk microscope using a 20× objective, 488-nm excitation, with images acquired every 2 s. After a 3-min baseline at 3 mM glucose, glucose was raised to 17 mM, and, at 18 min, KCl was added to 20 mM for Ca²⁺ imaging. Image analysis was performed in ImageJ using an in-house-made macro[25], available upon request.

For connectivity analysis, fluorescence traces and cell position of 15 control and 15 MTFP1-βKO islets, from 6 different animals, were extracted using ImageJ. The extracted calcium traced were analysed in MATLAB® as previously described[71], with minor modifications. Briefly, traces were smoothed by average and fold over baseline were calculated and used to produce raster plots[72]. Data were then binarized by identifying the minimum in a moving window representing 20% of the recording. Values deviating 5% from the local minimum were considered active. The second phase was identified as the activity following the first peak in islet and before the addition of KCl (typically 3 minutes after the onset of the increase in Ca^2+^, i.e. between the minutes 6 and 16 of the recording). Because the KO model did not display this first peak, the first phase was arbitrarily defined as the first 100 frames (200 s) following glucose addition. Coactivity was calculated on the second phase only as per [71], using the following formula:

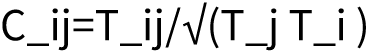

where Tij represents the time where two cells are contemporaneously active, while Tj and Ti is the total time where cell “i” and cell “j” are independently active. To identify whether a connection between two cell was significant a directional t-test was conducted on permutated data, where the binarized activity of each cell was randomly permutated 10,000 times and coactivity was recalculated at the end of each permutation. Significance was set at 2 standard deviations above the means (i.e. p=0.01)[73]. Connectivity was plotted using the MATLAB package wgplot. Dots represent cells while the connecting lines represent the coactivity value, ranging from red (high) to dark blue (low). The synchronicity between cells was plotted using the Bonferroni-corrected Pearson’s r obtained by correlation of the smoothed calcium traces. Total area under the curve was calculated using the trapezoid rule. Finally, peak width, frequency and duty cycle (i.e. percentage of total time spent active) was calculated for each cell. If a duty cycle resulted in 0, the cell was considered inactive.

### Tetramethylrhodamine ethyle ester (TMRE) imaging of mitochondrial membrane potential

Mitochondrial membrane potential (ΔΨm) was monitored in intact islets incubated with 10 nM TMRE in KRBH with 3 mM glucose and 0.1% fatty acid–free BSA for 45 min at 37°C. Islets were then transferred to bottom-glass chambers containing 2 nM TMRE, 3mM glucose in BSA-free KRBH. Imaging was performed using a 20x objective, 561-nm excitation, with images acquired every 2 s. After 3 min at 3 mM glucose, glucose was raised to 17 mM; at 16 min, 1 µM FCCP was added to collapse ΔΨm. Fluorescence traces were analysed as for Ca²⁺ and ATP imaging.

### Immunohistochemistry

Slides for immunohistochemistry were prepared from isolated formalin-fixed, paraffin-embedded pancreata and visualized as previously detailed[74]. For β-cell mass and proliferation measurements, slides were blotted with the following primary antibodies: anti-insulin (IR00261-2, 1:5; Agilent Techologies) and anti-glucagon (g2654, 1:100; Sigma) and/or anti-Ki67 (9129, 1:1000 Cell Signalling) antibodies, incubated with DAPI (Sigma-Aldrich) and mounted. To quantify apoptosis, pancreatic sections were, before mounting, submitted to terminal deoxynucleotidyl transferase dUTP nick-end labeling (TUNEL) assay using a DeadEnd Fluorometric TUNEL system kit (Promega) according to the manufacturer’s instructions. Slides were visualized with an Axiovert 200M microscope (Zeiss). ImageJ software was used to quantify the area positively stained for insulin and glucagon and the number of Insulin-, Ki67-, TUNEL- and DAPI-positive cells of all visible islets. For β-cell mass, we calculated the percentage of pancreatic surface that was positive, as measured in whole pancreas sections separated by at least 25 μm in the z-axis. For β-to-α cell mass we calculated the ratio of insulin to glucagon positive area in all visible islets. Ki-67- and TUNEL-positive β-cells were counted manually and expressed as a percentage of insulin-positive cells.

### Transmission electron microscopy and morphometric analysis

Transmission electron microscopy (TEM) of islets was performed as previously described[25]. Briefly, islets were fixed in 2% PFA + 2% glutaraldehyde in 0.1 M cacodylate buffer, washed twice with 0.1M cacodylate and stained with 2% osmium tetroxide for 1 h at 4°C, followed by 1% tannic acid. Samples were further washed in 0.05M cacodylate, dehydrated in graded ethanol and embedded in Epon resin. Ultrathin sections were cut with a diamond knife (DiATOME) in a Leica Ultracut UCT 6 ultramicrotome and imaged on a T12 Spirit TEM.

Cristae morphology was scored blindly using a semi-quantitative scale (0–4; 0— no sharply defined crista, 1— >50% of mitochondrial area without cristae, 2>greater than 25% of mitochondrial area without cristae, 3— > 75% of area crista, but irregular, 4—>75% area crista, mainly regular) and quantitative morphometric parameters were obtained in ImageJ, including cristae area, cristae volume density (cristae area/mitochondrial area), cristae surface density (cristae area/β-cell area), mitochondrial area fraction, form factor (perimeter²/4π·area) and circularity (4π·area/perimeter²), based on [75]. ER–mitochondria contact sites (MERCS) were defined as regions where ER and mitochondrial membranes were within 2–20 nm; the percentage of mitochondrial perimeter in contact with ER and the mean ER–mitochondria distance were manually drawn and quantified in ImageJ. Insulin granules were classified as mature, immature or empty/uncrystallised and counted per β-cell, blinded.

### RNA sequencing and analysis

RNA was extracted from 100-250 islets per mouse with TRIzol followed by TURBO DNase treatment (Thermo Fisher). RNA was further purified by acidic phenol:chloroform extraction and ethanol precipitation and quantity and integrity determined with an Agilent Bioanalyzer. Library preparation and sequencing was performed at the Imperial BRC Genomics Facility. Libraries were prepared from 400-1000ng RNA following mRNA enrichmend with a NEBNext Poly(A) mRNA Magnetic Isolation Kit (NEB), using a NEBNext Ultra II Directional RNA Library Prep Kit for Illumina (NEB) and Universal i5 and i7 primers following manufacturer’s instructions. Sequencing was performed on a HiSeq4000 using 75 bp paired end reads according to Illumina specifications. FastQC was used to assess the quality of the reads (∼50-90M/sample) that were then mapped to the mouse transcriptome (GRCm39) using Salmon v1.10.2[76]. Differential expression analysis was performed with the R package DESeq2 v1.46.0[77]. Genes with very low counts (average < 200) were excluded. Differentially expressed genes were defined by adjusted p-value (padj) < 0.1. Functional enrichment of differentially expressed transcripts was assessed using DAVID Gene Ontology analyses[78].

### Statistical analysis

Statistical significance was evaluated with GraphPad Prism 10.6.1 as indicated in the Figure legends. All data are shown as means ± SEM. p < 0.05 was considered statistically significant unless otherwise indicated.

## Acknowledgements

SS was supported by a Diabetes UK PhD studentship awarded to AMS (21/0006358). AMS and AMS’s lab (SS, RA, ZW, CZ, MP, MY, AM and BB) was also supported by a Project Grant (MR/X009912/1) by the Medical Research Council (MRC, UK) and a Project Grant (22/0006450) by the Diabetes UK. Human pancreatic islets were provided by the NIDDK-funded Integrated Islet Distribution Program (IIDP) (RRID:SCR_014387) at City of Hope, NIH Grant # U24DK098085 and the JDRF-funded IIDP Islet Award Initiative. A European Partners Fund from Imperial College London was awarded to AMS to import floxed MTFP1 mice from TW. G.A.R. was supported by a Wellcome Trust Investigator Award (WT212625/Z/18/Z), MRC Programme grant (MR/R022259/1), Diabetes UK (BDA 16/0005485) and NIH-NIDDK (R01DK135268, 1R01DK139630-01A1) project grants, a CIHR-JDRF Team grant (CIHR-IRSC TDP-186358 and JDRF 4-SRA-2023-1182-S-N), CRCHUM start-up funds, and Innovation Canada John R. Evans Leader Awards (CFI 42649, 46539). GO was supported by a Fonds de Recherche du Quebec en Santé Fellowship (#333390). TR was supported by MRC project grants (MR/N009371/1 and MR/T028637/1) and BBSRC project grant (BB/S008284/1). The Facility for Imaging by Light Microscopy (FILM) at Imperial College London is, in part, supported by funding from the Wellcome Trust (grant 104931/Z/14/Z) and BBSRC (grant BB/L015129/1). Tissue paraffinisation and sectioning were performed by the Imperial College Research Histology Facility. RNA sequencing data analyses were conducted on the Imperial College Research Computing Service, DOI: 10.14469/hpc/2232.

## Author contributions

SS & RA performed most experiments, designed research studies and obtained and analysed data; ZW, CZ, MP, MY, AM and BB generated experimental tools and performed research studies (i.e. CRISPR/Cas cells, virus, TEM, β-cell mass and apoptosis); GO performed connectivity analysis, TW provided MTFP1 plasmids and floxed mice. GR and TR contributed to the study design, with reagents and data interpretation. AMS conceived the study, designed experiments, analysed data, and wrote the paper; All authors read, provided feedback and approved the manuscript. AMS is the guarantor of this work and, as such, had full access to all the data in the study and takes responsibility for the integrity of the data.

## Conflicts of interest

GAR has received grant funding from, and is a consultant for, Sun Pharmaceuticals Inc. No other potential conflicts of interest relevant to this article were reported.

---

**Supplemental Figure 1.**
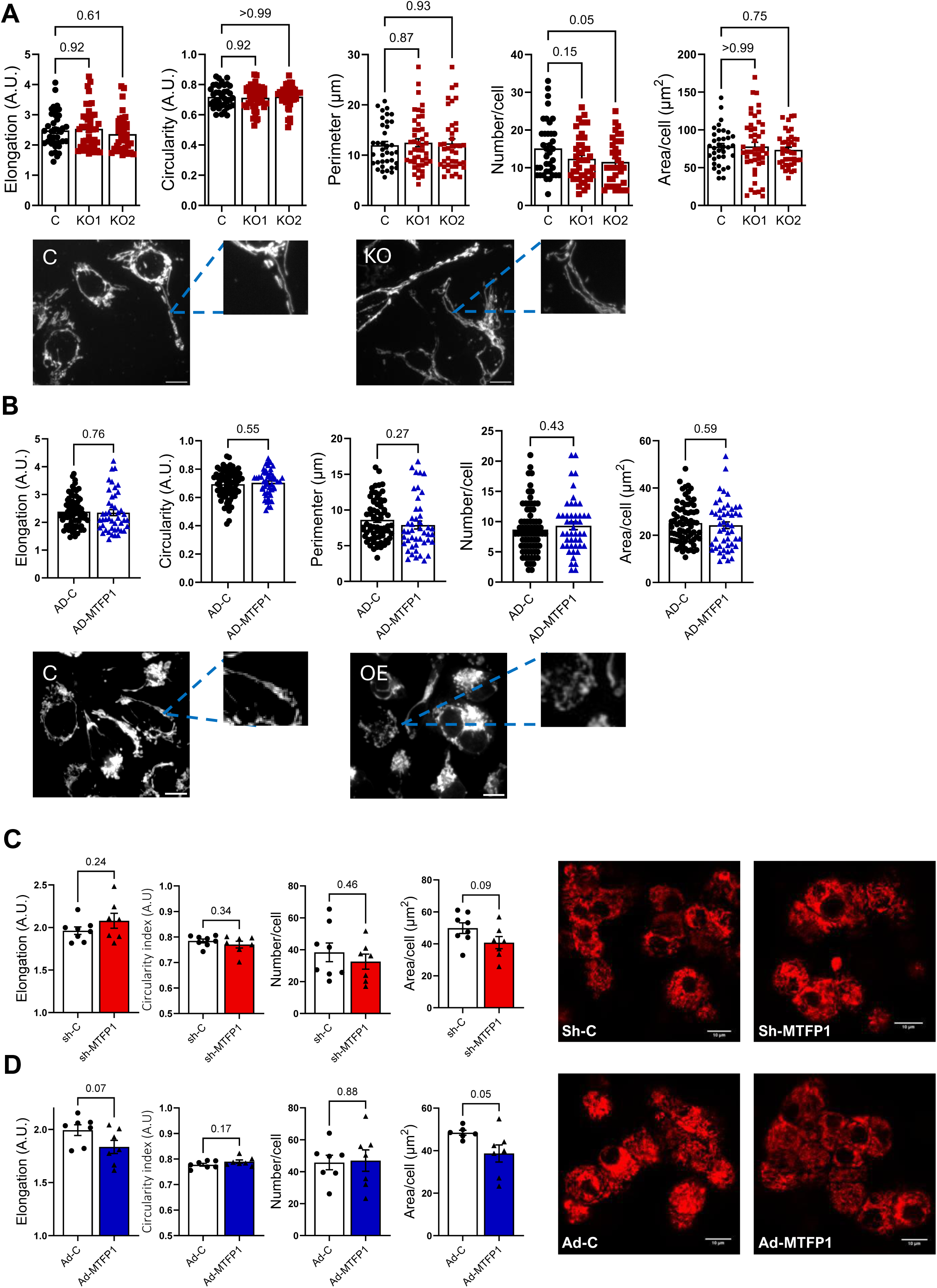
Loss- and gain-of MTFP1 function doesn’t affect mitochondrial elongation in EndoC-βH3 cells and human islets. **A)** MTFP1-KO1 (KO1) and MTFP1-KO2 (KO2) cells were generated with CRISPR/Cas9 targeting MTFP1. CRISPR/Cas9 EndoCβ-H3 populations were obtained via three independent lentiviral infections. **B)** EndoCβ-H3 cells Cells were infected with adenovirus expressing HA-tagged MTFP1 and GFP (AD-MTFP1) or GFP only (AD-C, controls). EndoCβ-H3 were treated with tamoxifen for 21-42 days. **C,D)** Human islets were infected with lentivirus carrying an shRNA targeting MTFP1 (sh-MTFP1, KD) or non-targeting (sh-C, C) at 5 MOI **(C)** or with adenovirus expressing HA-tagged MTFP1 and GFP (AD-MTFP1) or GFP only (AD-C, controls) at 10 MOI **(D)** for 72h. Mitochondrial network was stained with Mitotracker-green (A) or mitotracker-red (B, C, D) in live cells/islets and imaged using confocal microscopy. ImageJ was used to quantify individual mitochondrial length (elongation), circularity, perimeter, number of mitochondria per cell area imaged, and total mitochondria area per cell imaged. Representative images are shown on the bottom (A,B) or right-hand-side (C,D) panels. Scale bar= 10μm. Each dot represents an independent cell image (A,B) or average cell values in an independent islet (C,D), obtained in 3 separate experiments (n=3) or from two independent donots (C,D). Error bars represent SEM, one-way ANOVA with repeated measures and Dunnet multiple comparison test (A) and unpaired Student’s t test (B, C, D).

**Supplemental Figure 2.**
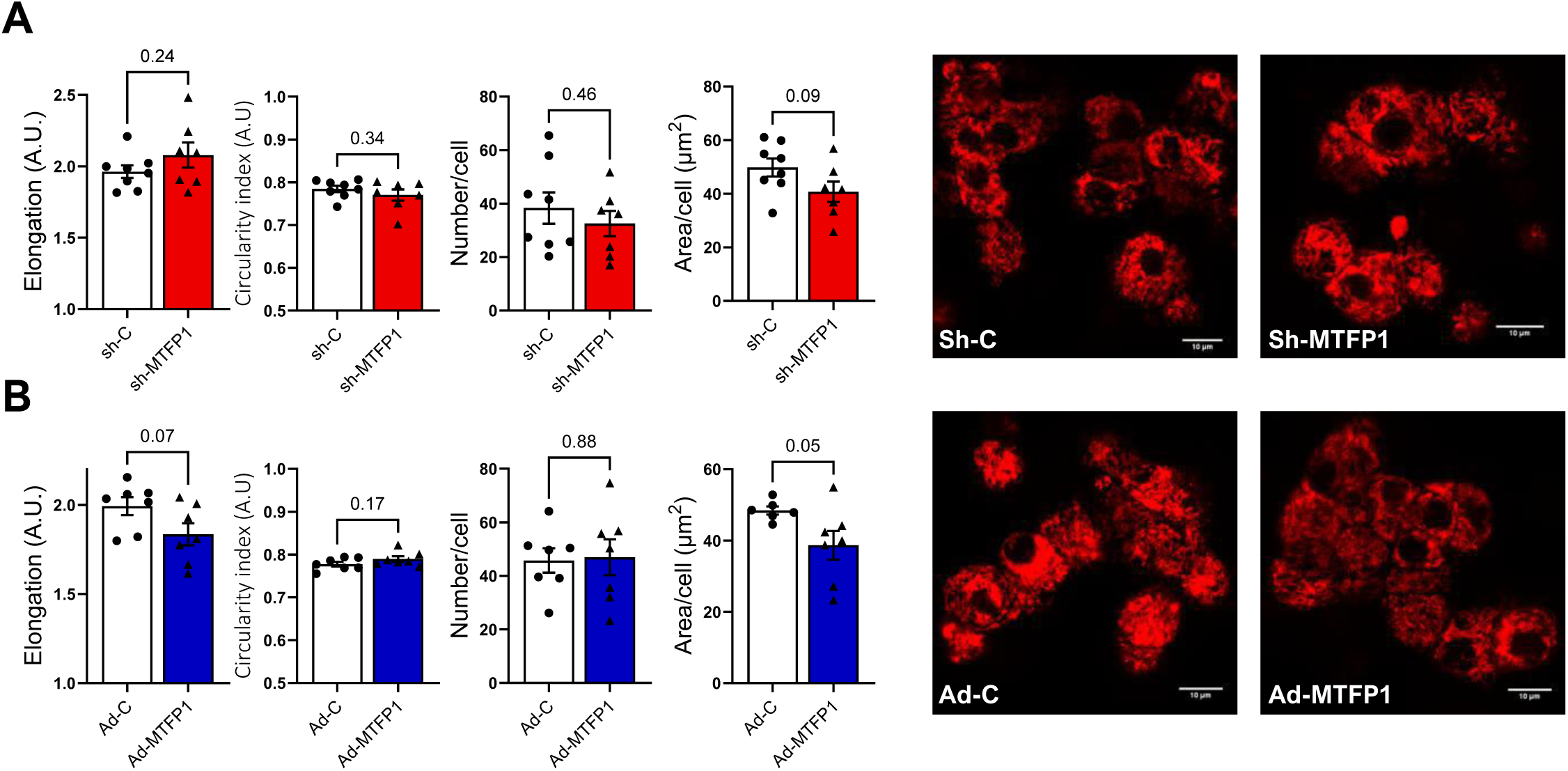
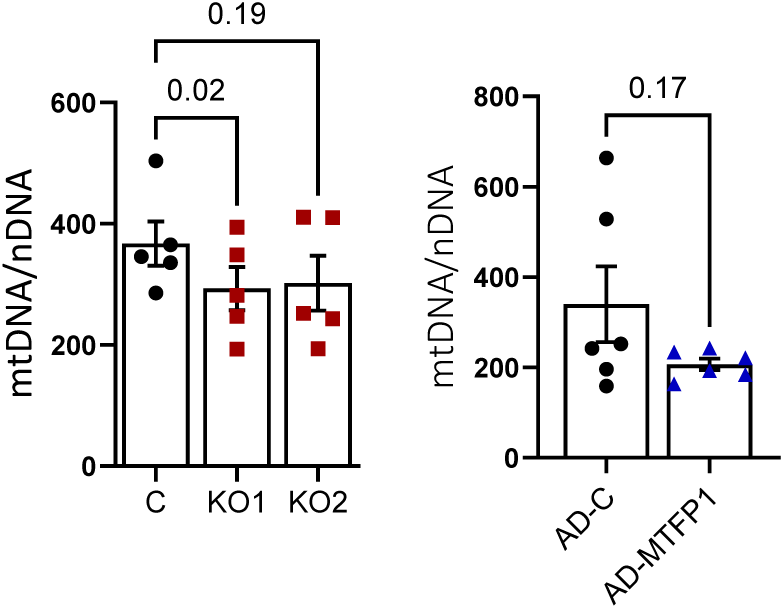
Effect of loss- and gain-of MTFP1 function in mitochondrial DNA content. **A)** MTFP1-KO1 (KO1) and MTFP1-KO2 (KO2) cells were generated with CRISPR/Cas9 targeting MTFP1. CRISPR/Cas9 EndoCβ-H3 populations were obtained via three independent lentiviral infections. EndoCβ-H3 were treated with tamoxifen for 21-42 days. **B)** Cells were infected with adenovirus expressing HA-tagged MTFP1 and GFP (AD-MTFP1) or GFP only (AD-C, controls) at 2 MOI for 48h preceding the experiments. Mitochondrial DNA copy number was calculated as the ratio of the mitochondrial encoded gene *mt-ND1* to the nuclear *B2G*. Each dot represents an independent experiment consisting of cells harvested on separate occasions. Error bars represent SEM, one-way ANOVA with repeated measures and Dunnet multiple comparisons test (A), and paired Student t test (B).

**Supplemental Figure 3.**
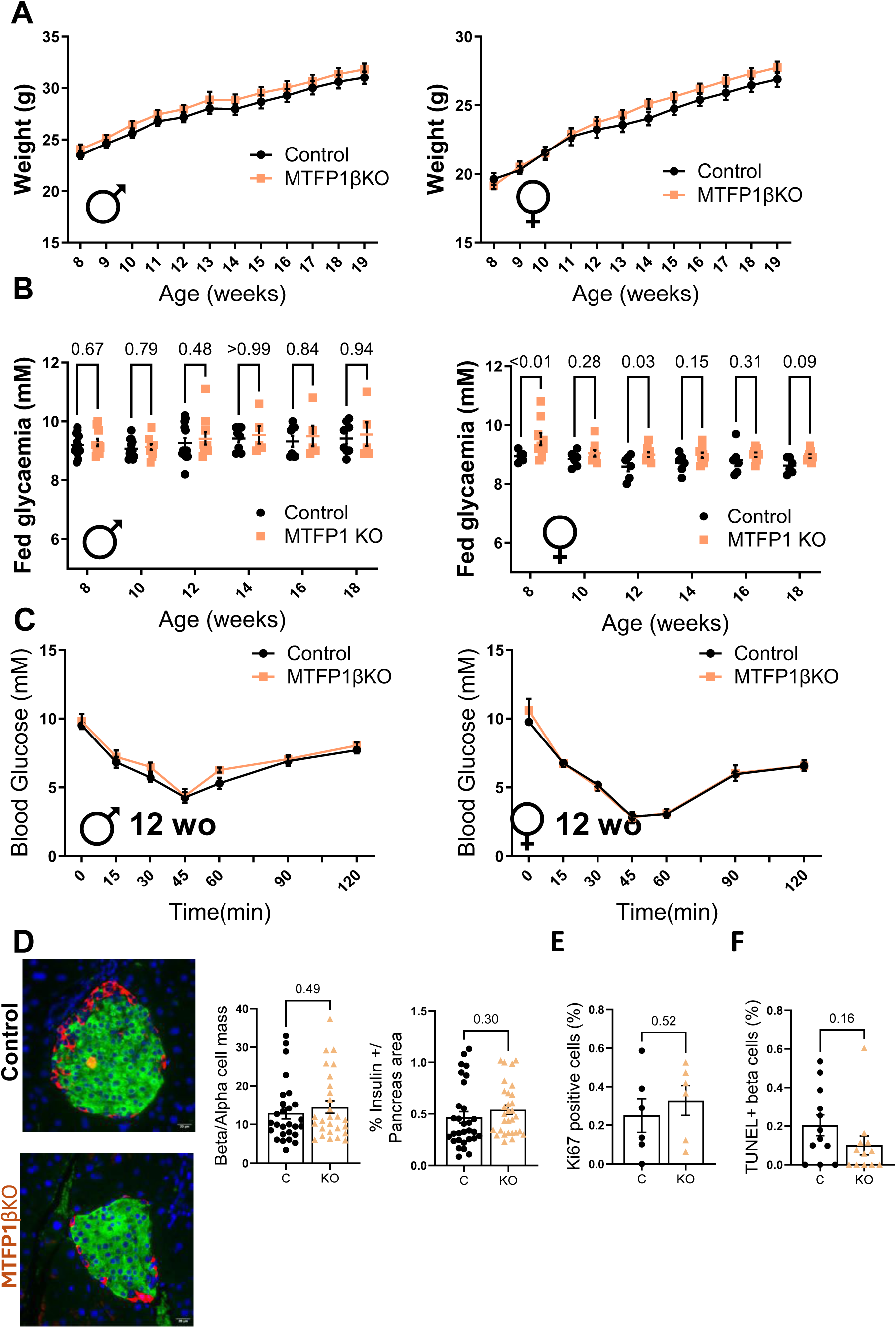
β-cell-selective elimination of MTFP1 does not alter body weight, randomly-fed glycaemia, insulin tolerance and β-cell mass. **A)** Body weight and **B)** randomly-fed glycaemia of male and female MTFP1βKO (MTFP1^f/f^, InsCre^+/−^) and littermate control (MTFP1^f/f^) mice monitored from the age of 8 weeks. n=5-13 mice/genotype. In (B) each dot represents a different mouse over time. **C)** Intraperitoneal Insulin tolerance test in 12-week old MTFP1βKO and littermate control male and female mice. n=5-8 mice/genotype (male)/2-3(female). **D)** Pancreata from 12 week – old male mice were fixed and subjected to immunocytochemical analysis and quantification for insulin and glucagon **(D)**, insulin and Ki67 and **(E)** insulin and TUNEL. Each dot represents one pancreatic section with n=3-5 mice/genotype. Error bars represent SEM, mixed effects analysis with Fisher’s LSD test (B) and unpaired t test with Welch’s correction (D-F).

**Supplemental Figure 4.**
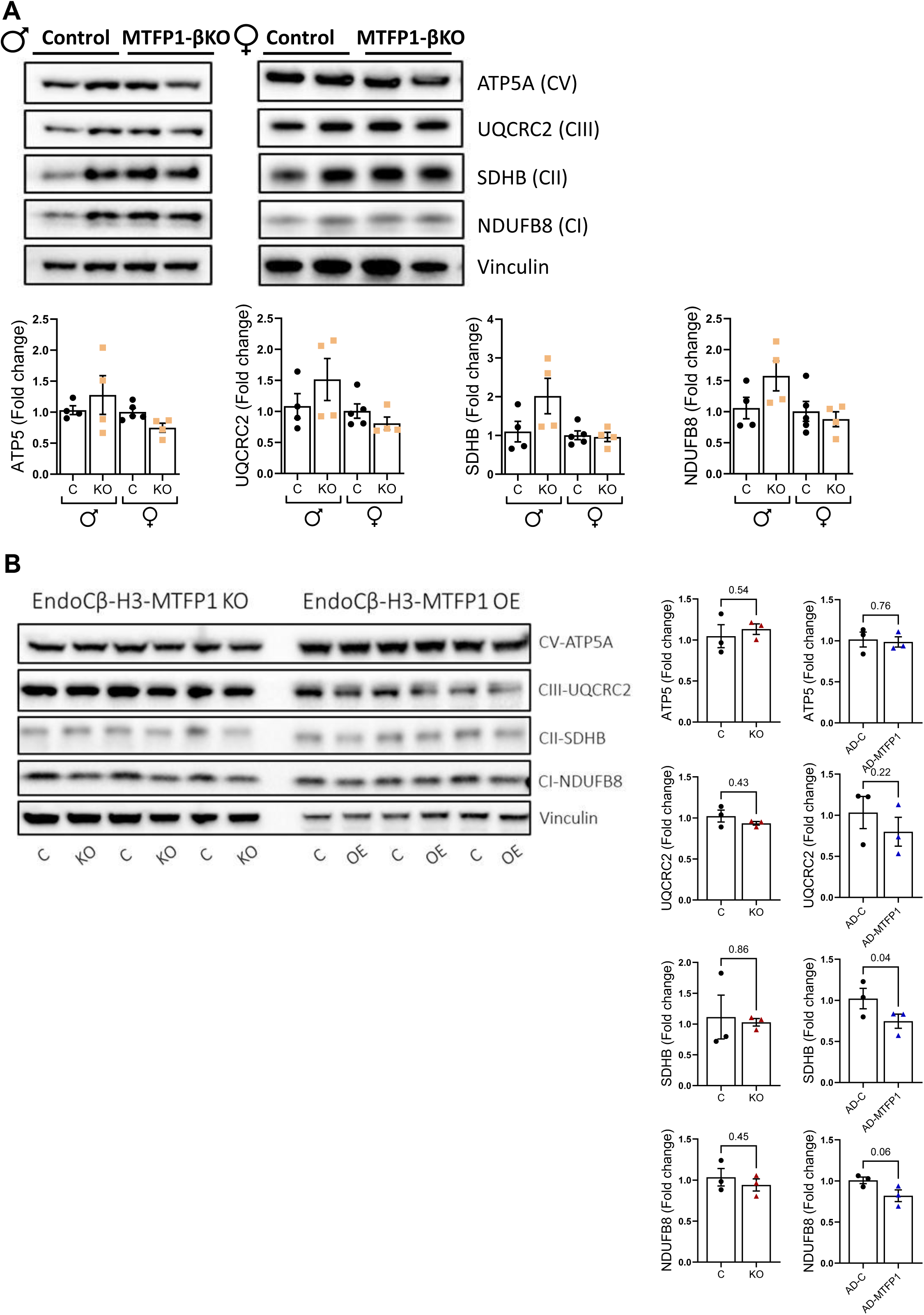
ETC complex I to V protein levels remain unchanged in MTFP1βKO islets. **A)** Representative Western blot (WB) showing expression of the indicated proteins in isolated islets from male (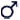) and female (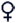) control (MTFP1^f/f^) and MTFP1βKO mice (MTFP1^f/f^, InsCre^+/−^). **B) Representative WB in** MTFP1-KO1 (KO) generated with CRISPR/Cas9 targeting MTFP1 versus non-targeting controls (C) and EndoCβ-H3 cells infected with adenovirus expressing HA-tagged MTFP1 and GFP (AD-MTFP1) or GFP only (AD-C, controls). Bar graphs show densitometry quantification normalized to vinculin. Each dot represents a different mouse, n=4-5 mice/genotype (A) or an independent experiment (B) n= 3-4. Error bars represent SEM, unpaired Student’s and Welch’s t tests (A) and paired Student’s t-test showed no significant (<0.05) differences.

**Supplemental Figure 5.**
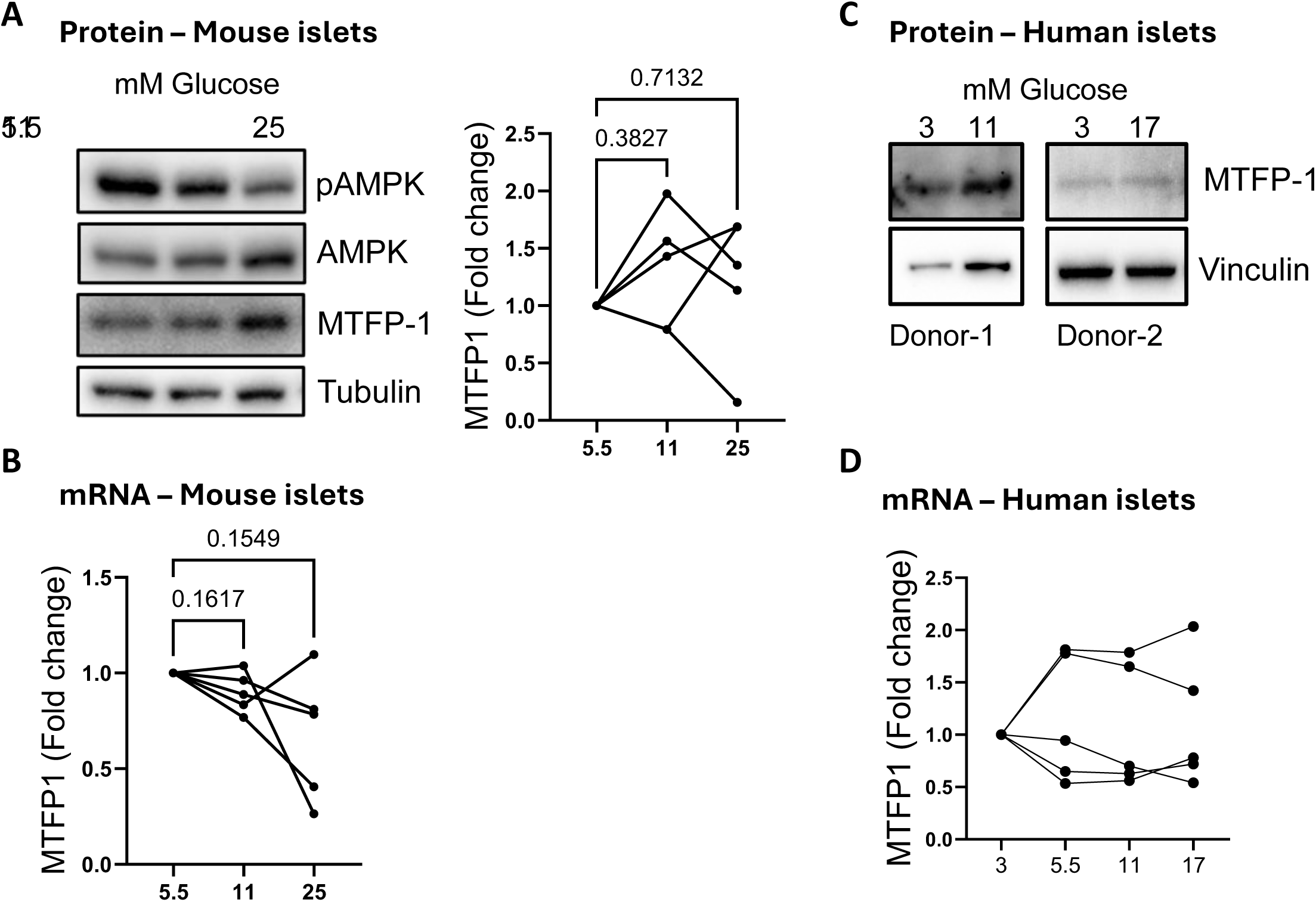
MTFP1 protein and mRNA levels remain unchanged in mouse and human islets cultured at high glucose concentrations. Islets from C57BL6/J male mice **(A, B)** and human donors **(C, D)** were cultured at different glucose concentrations for 48h. **A)** Representative Western blot showing protein levels of pThr172-AMPK, total AMPK, TMTFP1 and tubulin. Paired dot-plot shows densitometry quantification of mouse MTFP1 normalised by tubulin (n=5). **C)** Western blots (n=2, islets from two different donors). **B, D)** Quantitative RT-PCR measurements of MTFP1 normalized to Ppia (B) or RPLP0 (D). Data is presented relative to 5.5 mM (mouse) and 3mM (human) glucose. Each dot represent a different mouse (B) or donor (D). Samples used for D correspond to those prepared and described in Cheung et al (24). One-way ANOVA with repeated measures and Dunnet multiple comparisons test.

**Supplemental Figure 6.**
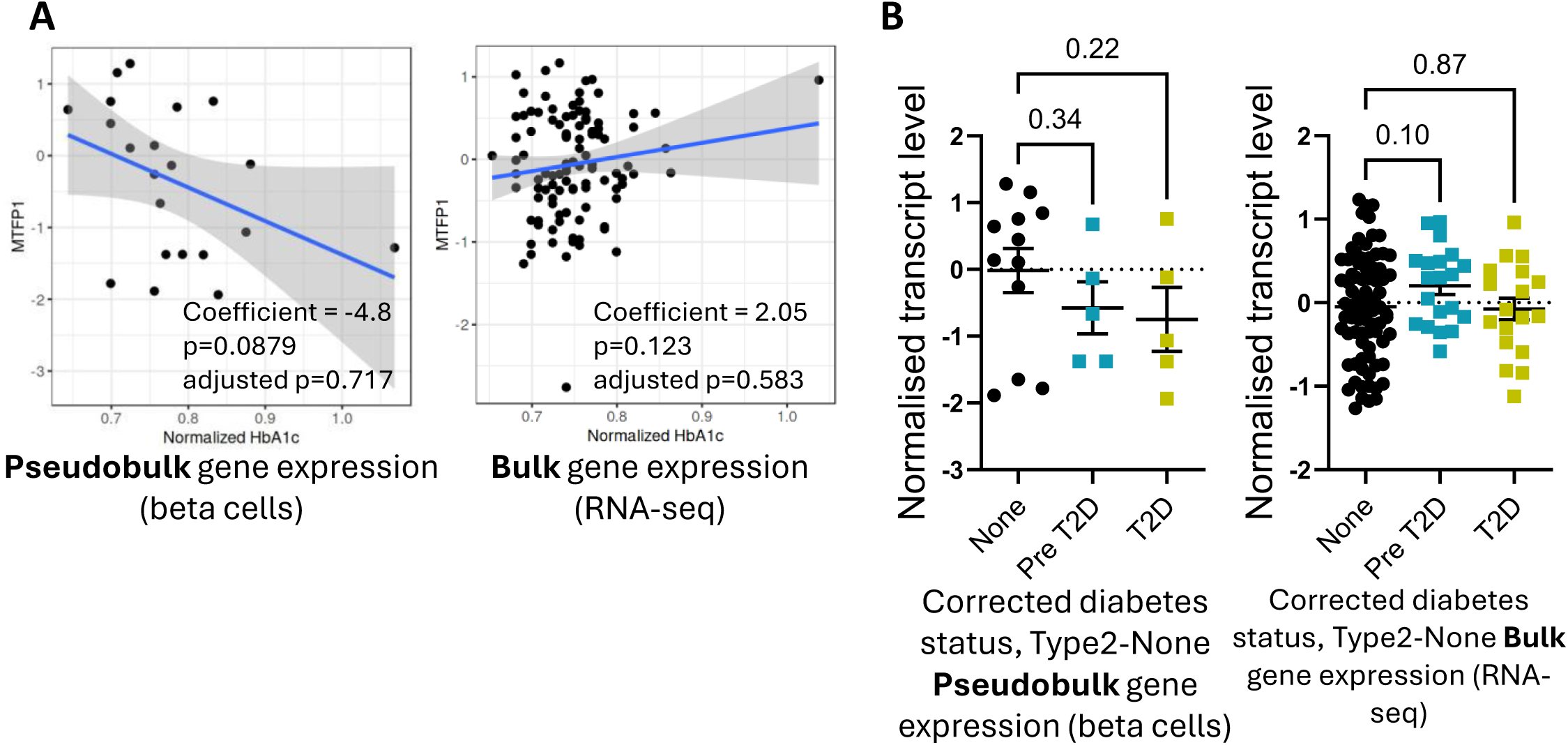
MTFP1 mRNA levels in β-cells inversely correlate with donor’s HbA1c and are lower in T2D than in healthy donors. Data downloaded from human islet project (https://www.humanislet.com). **A)** Correlation analysis of MTFP1 mRNA levels detected in beta cells by pseudobulk RNA-seq (left hand-side panel) and whole human islets by bulk RNA-seq (right hand-side panel) and HbA1c of the islet donors. **B)** MTFP1 mRNA levels detected in beta cells by pseudobulk RNA-seq (left hand-side panel) and whole human islets by bulk RNA-seq (right hand-side panel) from donors that lived with (Type2) or without (none) type 2 diabetes. Pre-T2D indicates pre-diabetes, defined as HbA1c of 5.7-6.5% and no clinical diagnosis of T2D). Error bars represent SEM, one-way ANOVA with Fisher’s LSD test.

## Notes

### Summary of Updates

We have increased the number of samples quantified for Mitotracker measurements and edited the figures accordingly. Overall content and conclusions of the manuscript were not affected by the results.

